# Label invariance: a guiding principle for ecological models

**DOI:** 10.1101/2025.07.15.664968

**Authors:** Theo Gibbs, Onofrio Mazzarisi, Lorenzo Fant, Ruby An, Jacopo Grilli, György Barabás, Chuliang Song

## Abstract

Ecological models, though diverse in form, are strengthened when they obey guiding principles. We formalize and advocate for a foundational principle we call “label invariance”, which says that a model’s dynamics must remain the same when identical individuals are arbitrarily grouped into distinct sub-populations. This principle is a necessary consequence of trait continuity—the observation that ecological interactions change continuously as organisms become more similar. Violation of label invariance often implies a hidden, intrinsic niche differentiation between species, which may obscure the mechanisms of biodiversity maintenance. We provide a general framework for constructing both deterministic and stochastic models that follow label invariance. We further demonstrate its utility as a complementary, non-statistical tool for empirical model selection. In sum, label invariance provides an important test for evaluating existing ecological models and a guide for developing new ones, promoting clarity in model assumptions from the outset.

## 1 What is label invariance

Universal principles in a scientific domain delineate what is possible, transcending the details of specific mechanisms (Lange, 2007; Maudlin, 2007). They are often the foundational ideas upon which many natural sciences are built: physics has the law of inertia and causality; chemistry has the conservation of mass and the law of definite proportions. Ecology, too, has its own set of principles that govern the dynamics of populations and communities (Turchin, 2013). Consider two such principles: First, because new organisms arise only from existing ones, models of population change are necessarily framed in terms of *per capita* growth rates. Second, since no environment offers limitless resources, these per capita growth rates inevitably become negative at high densities. These principles have actively guided the development of ecological models, and indeed, most contemporary formulations adhere to them. But models that obey just these two principles are not necessarily accurate representations of nature. The arbitrariness involved in the construction of new models suggests that ecology could benefit from formalizing additional model-building principles.

In this paper, we argue that *label invariance* is another principle for ecology. To understand this principle, consider a simplified ecological system comprising two species, *x* and *y*, where all individuals within a species are identical (Figure 1). Observer 1 correctly identifies these species and measures all parameters necessary for an ecological model. Observer 2, while also accurately measuring all relevant parameters, identifies the same system but subdivides species *x* into two subspecies, *x*_1_ and *x*_2_. Label invariance dictates that a valid ecological model should produce identical dynamics for both observers, as the underlying ecological reality is unchanged. Specifically, the total abundance of *x* (as perceived by Observer 1) must equal the combined abundance of *x*_1_ and *x*_2_ (as perceived by Observer 2), and the abundance of *y* must remain constant across both observers. In short, if there is no difference between species in ecological reality, our models should not predict one based on observer labeling.

**Figure 1.**
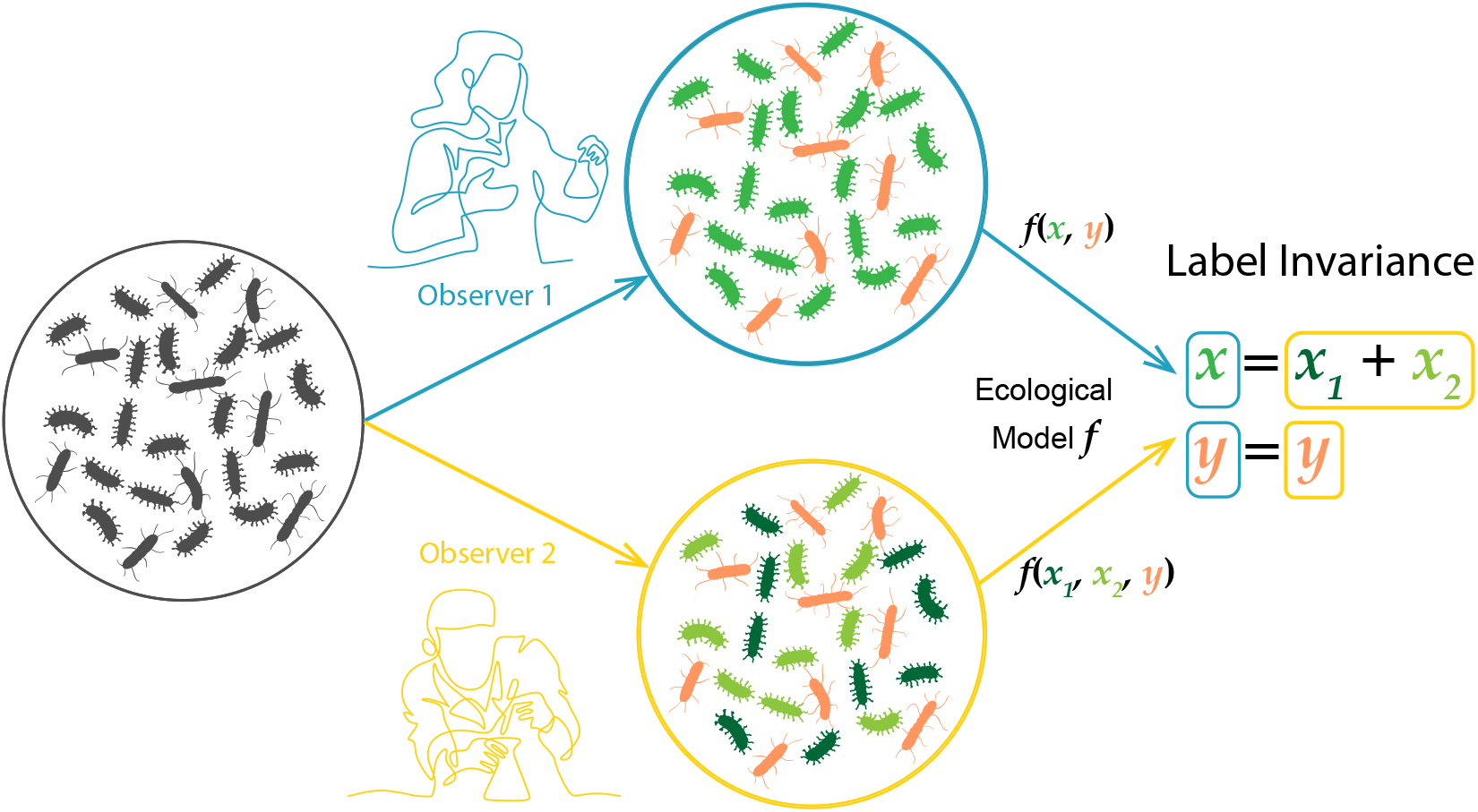
Illustration of label invariance. A single ecological community (left) is classified differently by two observers. Observer 1 identifies two species, *x* (green) and *y* (orange). Observer 2, viewing the same community, also identifies species *y* but chooses to subdivide the functionally identical individuals of species *x* into two arbitrary subgroups, *x* _1_ (dark green) and *x* _2_ (light green). The principle of label invariance requires that a valid ecological model, *f*, must produce consistent dynamics regardless of the observer’s labeling scheme. This means the model’s output must satisfy the conditions that the total abundance of the subdivided groups equals the single group (*x* = *x*_1_ + *x*_2_) and the abundance of the other species remains the same (*y* = *y*). In essence, a model’s predictions should not depend on the arbitrary labels we assign to them.

The principle we call ‘label invariance’ builds on a persistent, yet fragmented, lineage of similar ideas in the ecological literature. Since the early 1990s to the present (Ansmann & Bollenbach, 2021; Moisset de Espanes *et al*., 2021), this fundamental concept has repeatedly surfaced (see Section 8 for a more comprehensive historical review), albeit under different names such as ‘clone-consistency’, ‘common sense condition’, or ‘invariance under relabeling’. Our chosen term is a concise version of this last one. Past work has used mathematical techniques to classify all label invariant models, but these insights have arguably not yet permeated the broader ecological modeling community. We argue that this oversight may stem from a lack of connection between the mathematical definition of label invariance and its ecological underpinning. Similarly, past work has not explored the consequences of the violation of label invariance. This paper aims to bridge these gaps by rendering key mathematical ideas from previous work more accessible and by presenting new analytical insights.

We begin with an example of how to determine whether a model is label invariant, specifically considering the canonical Lotka–Volterra model. We then argue that the ecological underpinning of label invariance is the continuous nature of ecological traits and interactions—a foundation previously under-articulated. With this grounding, we critically examine the consequences of violating this principle, revealing what such violations imply in terms of niche differentiation. To address this, we offer guidance on formulating models that inherently respect label invariance for both deterministic and stochastic dynamics. We further demonstrate the practical utility of this principle in empirical contexts by showing how it aids in model selection when confronted with real-world data. After contextualizing these ideas in a brief historical overview of related concepts, we conclude by discussing the broader implications of adopting label invariance for the rigorous development of ecological theory.

### Box 1

Lokta – Volterra dynamics obey label invariance

To illustrate the mathematical basis of this principle, we show the Lotka–Volterra model, a cornerstone of theoretical ecology, is label invariant. For the sake of clarity, we will focus on a simplified scenario involving two competing species, *x* and *y* (a more generalized proof is provided in Appendix A; also see Moisset de Espanes *et al*. 2021). From the perspective of Observer 1 (Figure 2), the Lotka–Volterra dynamics unfold as follows:

**Figure 2.**
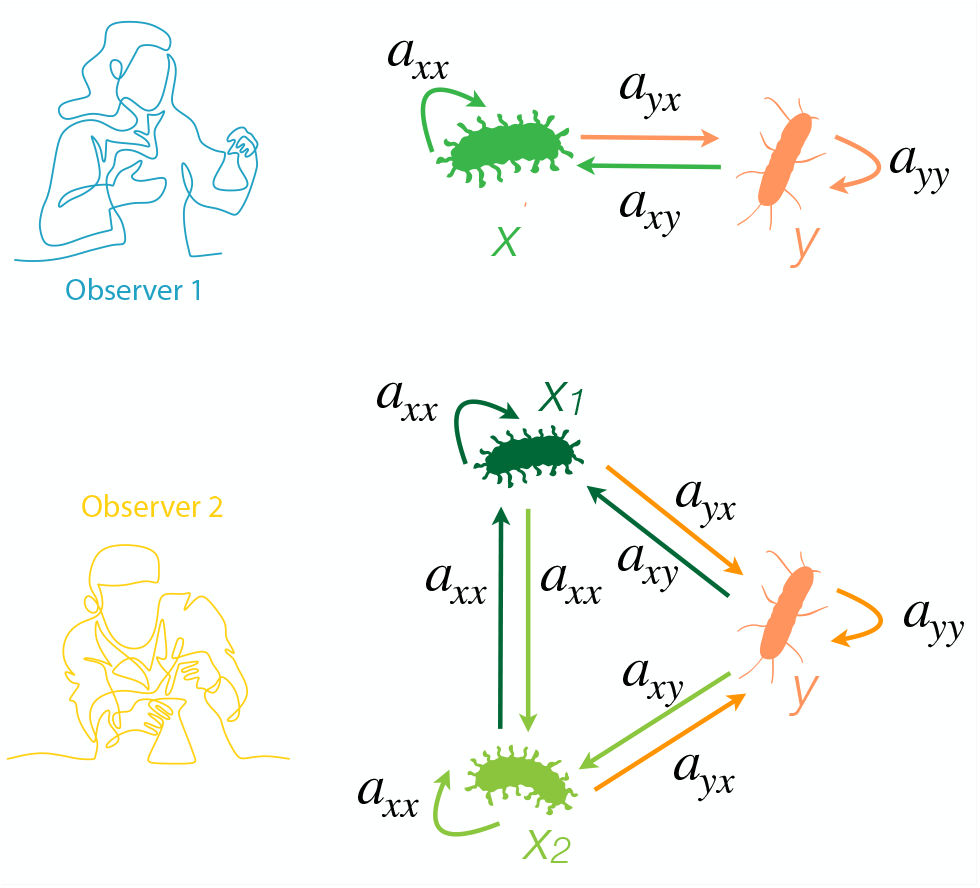
Lotka-Volterra model is label invariant. In Observer 1’s standard view (top), the effect of species x on itself (intraspecific competition) is defined by the coefficient *a*_*xx*_. When Observer 2 splits species x into two identical sub-species, *x*_1_ and *x*_2_ (bottom), the model’s consistency depends on a key logical step: the competitive interaction between the identical sub-species *x*_1_ and *x*_2_ must be the same as the interaction within the original species x. Therefore, the model assigns the same coefficient, *a*_*xx*_, to describe this interaction. Because the model’s structure ensures that interactions between identical individuals are treated consistently whether they are grouped together or split apart, the overall system dynamics remain unchanged, and the model is label-invariant.

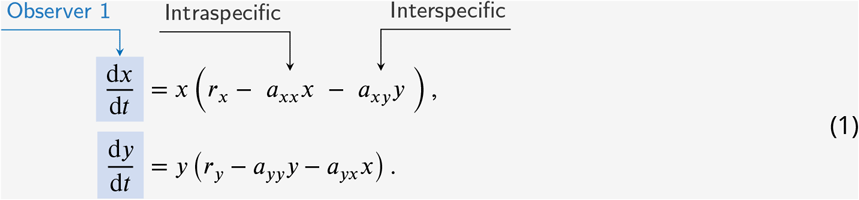

where *r*_*x*_ and *r*_*y*_ represent the intrinsic growth rates of the species, *a*_*xx*_ and *a*_*yy*_ represent intraspecific competition, and *a*_*xy*_ and *a*_*yx*_ represent interspecific competition.

Now, consider Observer 2, who observes the same ecological system but chooses to divide species

*x* into two subpopulations, *x*_1_ and *x*_2_. The Lotka–Volterra equations, from this perspective (Figure 2), transform into:

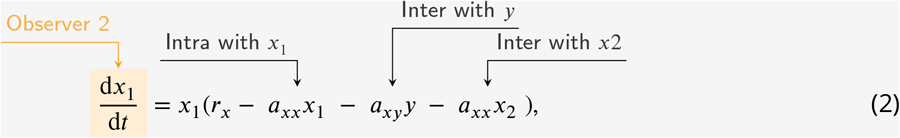

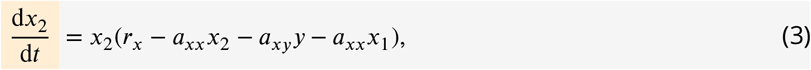

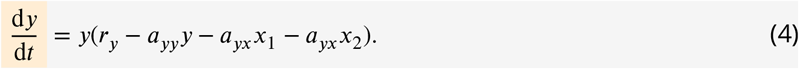

Since individuals in sub-species *x*_1_ and *x*_2_ are fundamentally identical, the interactions *between* them are equivalent in strength to the interactions *within* the original species *x*. This translates into the interspecific competition coefficient between *x*_1_ and *x*_2_ being equal to the original intraspecific competition coefficient (*a*_*xx*_).

The principle of label invariance demands that the dynamics of the system remain consistent, regardless of whether we adopt the perspective of Observer 1 or Observer 2. To verify this, let’s sum the equations for *x*_1_ and *x*_2_ and observe how *y* changes under Observer 2’s viewpoint:

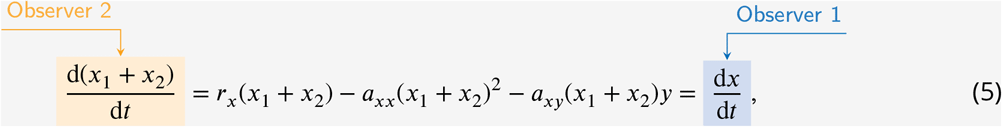

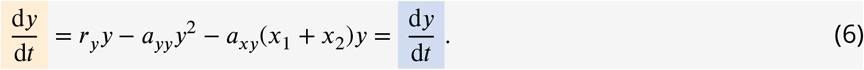

We find that the dynamics governing the total abundance of *x* as seen by Observer 1 perfectly mirror the combined dynamics of *x*_1_ and *x*_2_ as perceived by Observer 2. Moreover, the dynamics of *y* remain unchanged across both perspectives. This agreement confirms that the Lotka–Volterra model adheres to the principle of label invariance. The arbitrary reclassification by Observer 2 did not alter the model’s fundamental ecological predictions.

There is an even easier way of demonstrating label invariance. For any closed ecological system without immigration, the dynamics of its species can be written in terms of per capita growth rates: d*x*_*i*_ /d_*t*_ = *x*_*i*_***r***_*i*_(*x*_1_, *x*_2_, …),, where *x*_*i*_ is species *i*’s population density and ***r***_*i*_(*x*_1_, *x*_2_, …) is its per capita growth rate. The per capita growth rate pertains to individuals, and label invariance dictates that identical individuals, at any given time, must have identical (or identically distributed in a stochastic setting) per capita growth rates, regardless of how we partition them into various subspecies. A model is therefore observer-invariant if the per capita growth rates are equal across otherwise identical subspecies. In the context of Eqs. 2-3, the parenthesized terms on the right hand sides are the appropriate per capita growth rates – and these are manifestly equal.

## 2 The ecological underpinnings of label invariance

Label invariance holds because the dynamics of an ecosystem are dictated by what organisms *do*, not what we *call* them. It is the ecological counterpart of the proverbial duck test: if individuals are indistinguishable based on their ecologically relevant traits, then the decision to classify them as a single species or as multiple distinct (but identical) subspecies should not influence the underlying ecological processes at play. This principle is independent of whether two individuals are truly identical, or identical only in all the ways they can interact with their environment. Further, we argue that label invariance is a consequence of the continuous nature of ecological processes. Organisms rarely fall into perfectly discrete bins based on their functional roles. In most real ecological situations, continuously varying traits are the norm (beak morphology, metabolic efficiencies, tree heights), producing continuous variation in ecological similarity.

As a conceptual example, we can picture two distinct groups of organisms and imagine their ecologically relevant traits gradually converging until they become functionally indistinguishable (Figure 3). An ecologically sound model must mirror this smooth transition. Its predictions for their combined dynamics should change continuously as the groups approach identity, without a sudden shifts as they move from being *very similar* to *exactly alike*. When the two groups are functionally identical, the model should predict the exact same dynamics for the total population, regardless of whether we label it as one species or two (now identical) subspecies, because, at this endpoint, any distinction is purely based on nomenclature. Consequently, differences in a model’s predictions for identical subpopulations signal an artificial discontinuity.

**Figure 3.**
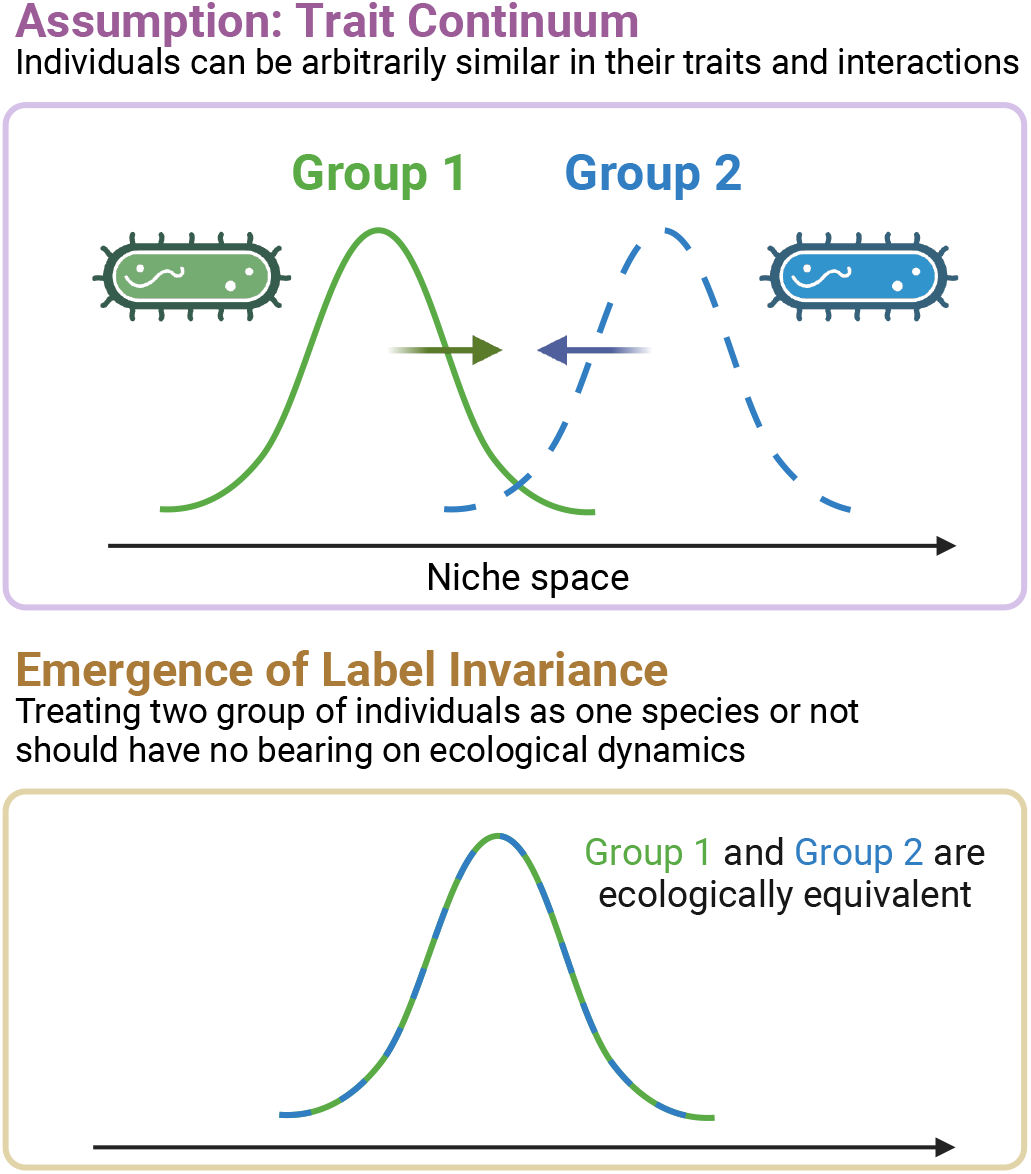
Label invariance is a logical consequence of trait continuity. This figure illustrates how the principle of label invariance emerges directly from the concept of trait continuity. The top panel depicts two distinct groups (Group 1 and Group 2) that can converge in niche space, becoming progressively more similar. An ecologically sound model must reflect this continuous transition, and the bottom panel shows the logical endpoint where the groups become ecologically identical and indistinguishable. At this point of perfect identity, the model’s predictions should not create an artificial “jump” or discontinuity. Therefore, the requirement that the model must produce the same dynamics whether the organisms are labeled as two identical sub-species or as one single species—which is the principle of label invariance—is a necessary consequence of this continuity.

This property—that models reflect the gradual convergence of dynamics as organisms become interchangeable—has been expressed mathematically through the principle of trait continuity (Meszéna, 2005; Meszéna *et al*., 2005). It states that, if per capita growth rates are functions of underlying traits, then a marginal tweak in those traits should provoke only a marginal response in the growth rates. In Leibniz’s words, *Natura non facit saltus*—Nature does not make jumps. Appendix B provides a formal mathematical derivation based on functional analysis. This is where ecology differs from the indivisible units in physics or chemistry. An H_2_O molecule cannot be “almost H_2_O” yet still retain the essential functional properties of water in a chemical reaction. Thus, for models built upon such irreducibly discrete components, label invariance does not apply. Indeed, there are (rare) ecological situations that are truly discontinuous. For example, communities of oligonucleotide replicators (von Kiedrowski, 1986) serve as both genotype and phenotype (Cech, 2012), and since different types of replicators must differ from one another by a single nucleotide at the very least, we have a discrete, countable set of possible phenotypes (Meszéna & Szathmáry, 2001).

On the whole, trait continuity—the idea that similar phenotypes yield similar per capita growth rates—is common because it emerges from more mechanistic underpinnings of many ecological processes. In consumer-resource interactions, the overlap in resource utilization between consumers is often a continuous variable (Appendix C). In ecophysiology, species experience limitations from light or water availability along a continuum (Appendix D). Indeed, one can ask the question: what would happen if we took some species (say, the medium ground finch *Geospiza fortis*) and arranged somehow to suddenly increase the bill depth of each of its individuals by 1 *μ*m? The answer is that nothing in the world would even notice that a change has occurred (Meszéna, 2005; Barabás *et al*., 2013). Consequently, if we employ a phenomenological model that violates the principle of trait continuity, it could signal a deficiency in plausible mechanistic grounding, or underlie important assumptions regarding differences between the populations modeled.

## 3 Label invariance reveals hidden niche differentiation

Models that do not satisfy label invariance may implicitly assume some degree of irreducible niche differentiation, even when species verge on perfect similarity in their modeled traits. In this section, we explore how such implicit niche differences play a crucial role in models that address biodiversity limits and ecosystem stability. First, we consider models that challenge the principle of limiting similarity, a foundational concept that imposes a theoretical cap on biodiversity by predicting that competition will exclude species with excessive niche overlap (MacArthur & Levins, 1967; Gyllenberg & Meszéna, 2005; Meszéna *et al*., 2006; Barabás *et al*., 2012; Barabás *et al*., 2014). Second, we show that the hidden nature of niche differentiation can also play a crucial role in the long-standing diversity-stability paradox (May, 1972; McCann, 2000). Specifically, we analyze a sublinear growth model (Hatton *et al*., 2024) and show that hidden niche differentiation is associated with mechanisms unique to each population.

### 3.1 Emergent neutrality model

In the emergent neutrality model (Scheffer & Van Nes, 2006; Vergnon *et al*., 2012), species are characterized by their positions along a trait axis, which determines how those species interact with one another. For example, the species could be granivorous birds competing for food, with the measured trait being their beak size. More similar beak morphologies are suited to consuming more similar food, and so competition between species increases with their degree of trait similarity. Additionally, each species experiences chronic pressure from natural enemies whose dynamics are unmodeled. This pressure is species-specific: each species experiences a pressure that is independent of the pressure on the others.

The model reads

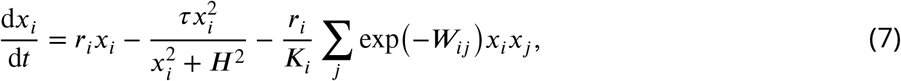

where *x*_*i*_, *r*_*i*_ and *K*_*i*_ are the population density, intrinsic growth rate, and carrying capacity of species *i, τ* and *H* are parameters of the type III functional response representing chronic pressure from the natural enemies, and *W*_*ij*_ is a unitless measure of the trait distance between species *i* and *j*. For the example of the bird community, *W*_*ij*_ would be proportional to the absolute difference of their beak sizes. The per capita growth rate of species *i* is obtained by dividing the right-hand side of Eq. 7 with *x*_*i*_:

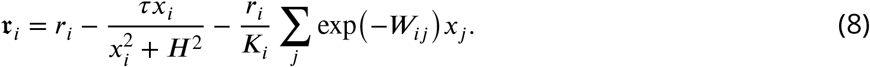

It has been shown that, in this model, very similar species can still coexist (Scheffer & Van Nes, 2006; Vergnon *et al*., 2012). This is true in the sense that *W*_*ij*_ ≈ 0 indeed does not preclude coexistence. However, the similarity between species is not entirely captured by their explicitly modeled trait values (beak size in the bird example). To see why, we first show that the per capita growth rates are not equal even when species *i* and *j* are identical in all their parameters, including beak size (*W*_*ij*_ = 0) and the intrinsic growth rates *r*_1_ = *r*_2_ ≡ *r*:

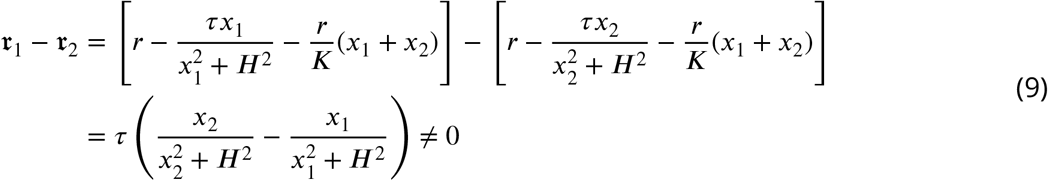

(since exp (− *W*_*ij*_) = exp(0) = 1, these factors were not written out). The reason these are not equal is the term 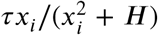 describing the chronic effect of the the unmodeled natural enemies. In short, this model is not label invariant. Each species experiences the corresponding negative density-dependent effects independently, regardless of how similar they are to other species (see Appendix E). This effectively assumes that there are unmodeled differences between the species which are irrelevant to their resource acquisition, but which can be used by the enemies to distinguish them (Barabás *et al*., 2013). In the bird example, species with identical beak morphology might possess different blood chemistry, which protects them from different sets of pathogens. Whatever the cause, there must be some unmodeled differences present to explain why this term is strictly species-specific. The fact that the model is not label invariant, therefore, reveals a tacit assumption of hidden trait differences, even between species that happen to be identical in the one trait that is explicitly modeled.

### 3.2 Sublinear growth model

A recently proposed sublinear growth model (Hatton *et al*., 2024) exhibits increased stability with higher species richness (May, 1972; McCann, 2000). This model modifies the classic Lotka–Volterra model by adding sublinear density dependence to the growth rate:

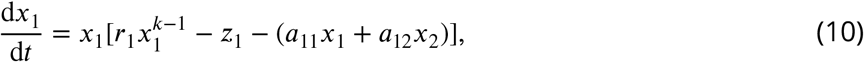

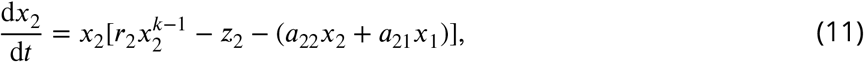

where *r*_1_ and *r*_2_ represent intrinsic growth rates, *z*_1_ and *z*_2_ are species decay rates, *a*_12_ and *a*_21_ represent interspecific competition and *a*_11_ and *a*_22_ intraspecific competition^1^, and the constant *<* 1 captures sublinear growth. The per capita growth rates *r* _1_ and *r* _2_ are given by the bracketed terms on the right hand sides of Eqs. 10-11.

However, these growth rates are not equal to one another when *x*_1_ and *x*_2_ purportedly represent identical populations—that is, when *r* _1_ = *r* _2_ ≡ *r, z*_1_ = *z*_2_ ≡ *z*, and *a* _*ij*_ ≡ *a* for any *i, j* ∈ 1, 2:

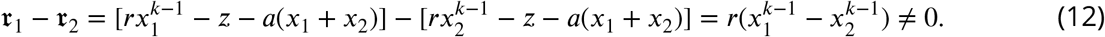

By employing an intuitive and broadly applicable definition of niche differentiation (Spaak & De Laender, 2020), we can show that two subpopulations with identical traits have an irreducible niche difference in this and more general models with the same diversity-stability properties (see Appendix E). This also implies that adding species to the system decreases niche overlap, even if they share the same traits as pre-existing ones.

The root of this mismatch is that we are not considering truly identical subpopulations. Instead, there is a hidden trait that is kept implicitly different between them. By casting the model in a label invariant form, we can reveal this hidden trait as the effect on the intrinsic growth rate from other species:

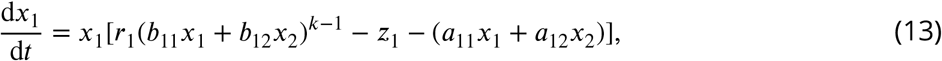

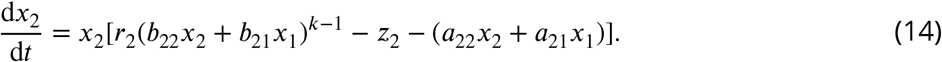

This effect is quantified by the matrix *b*. By increasing the number of parameters, we allow for other species to contribute to the sublinear density effects on growth. With the additional constraint that for identical *x*_1_ and *x*_2_, we have *b*_*ij*_ ≡ *b*, the per capita growth rates are now equal:

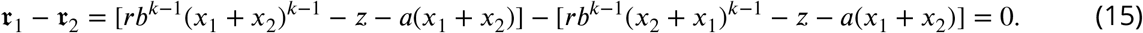

Two identical sub-populations, sharing the same set of parameters, will have complete niche overlap in this formulation. Importantly, to preserve the result in Hatton *et al*. (2024), the off-diagonal terms of the matrix *b* should be strictly zero (see Appendix F).

Through the lens of label invariance, we are able to highlight that only an increase in the number of species with an interaction mechanism that uniquely involves individuals within a population (in this case, sublinear self-regulation, guaranteed by the vanishing off-diagonal element of *b*), can stabilize the system. The within-species nature of the mechanism, however, is only a necessary condition; see papers by Hatton *et al*. (2024) and Mazzarisi & Smerlak (2024) for more details.

### 3.3 Label invariant formulations clarify mechanisms for coexistence

Violating label invariance is not inherently problematic. It can, however, allow for the emergence of hidden niche differentiation, which in turn weakens our control over the unstated assumptions embedded within a model, ultimately obscuring the true drivers of observed ecological patterns. In both the emergent neutrality and sublinear models, there is an interaction mechanism that uniquely applies to individuals within a population. This interaction mechanism creates both the hidden niche differentiation which leads to novel ecological patterns and the violation of label invariance through a discontinuous relationship between per-capita growth rates and traits values. Including interaction mechanisms that are unique to each population in our models may have no important consequences in some ecological situations, but when analyzing more generic diversity properties, it narrows the applicability and validity of these models to communities where each species differs profoundly along at least one relevant ecological axis.

## 4 How to build label invariant models

How does one construct models that are label invariant? The multispecies Lotka–Volterra model, as we have shown, label invariant. But it is not the only possible formulation, nor is it always the most appropriate for every ecological question. In this section, we describe the general principles that ensure label invariance, particularly as we move towards models incorporating more complex mechanisms. Illustrating this challenge, many models in the ecological literature are not label invariant – a survey of ten ecological models by Moisset de Espanes *et al*. (2021) found only one formulation (Bastolla *et al*., 2009) that passes this criterion.

### 4.1 Motivating example: how to model saturating functional responses

The subtleties of ensuring label invariance become particularly apparent when modeling phenomena like saturating functional responses. Consider a system with two resources *x*_1_ and *x*_2_. Three plausible formulations are:

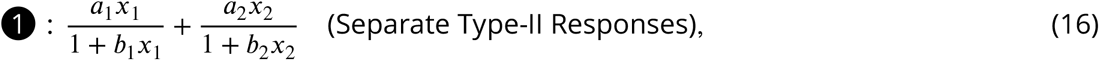

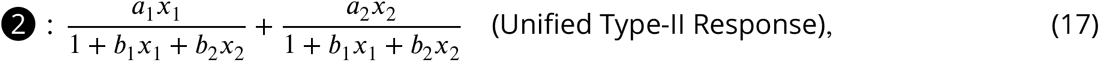

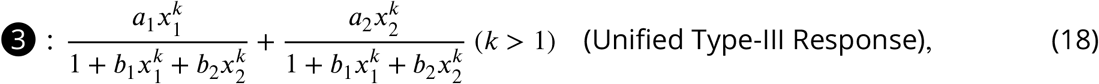

where *a* terms are attack rate/capture efficiencies, and *b* terms are handling times—how long it takes to process a food item, which causes saturation. In the absence of the label invariance criterion, one might argue that all three choices adequately capture the concept of a saturating effect. Indeed, each of these functional responses has appeared in the ecological modeling literature (e.g., Qian & Akçay 2020 and Aguadé-Gorgorió *et al*. 2024 used 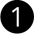; Thébault & Fontaine 2010 used 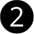; Ryabchenko *et al*. 1997 used 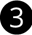). However, only formulation 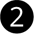 is label invariant (Arditi & Ginzburg, 2012; Morozov & Petrovskii, 2013).

The crucial difference is how these models describe the predator’s processing bottleneck—what limits its intake when food is abundant. If we take a fixed quantity of resource *x*_1_ and divide it into *n* equally abundant but perfectly identical subcategories, the predator cannot distinguish between them; they are all functionally the same resource from its point of view.

Formulation 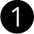(Separate Type-II) treats the saturation for each listed resource category as independent, effectively giving the predator multiple independent stomachs. As *n* increases (or as we arbitrarily sub-divide the identical resource), this model predicts that the predator’s total intake rate of *x*_1_ will increase. In contrast, Formulation 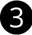(Unified Type-III), with its nonlinear aggregation (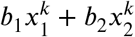 where *k >* 1), leads to the opposite but equally unrealistic outcome. When *n* increases, this model predicts that the predator’s total intake rate of *x*_1_ will decrease. Neither scenario reflects biological reality; the predator’s digestive system doesn’t change based on how we name the resources.

In the observer-invariant Formulation 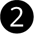(Unified Type-II), a single, shared constraint limits the processing of all food items, analogous to having one single stomach. This model predicts that the total intake rate of *x*_1_ remains unchanged, because the influences of different resources (or subdivisions of the same resource) are combined *linearly* within both the numerator and the denominator.

### 4.2 Linear pathways guarantee label invariance

The linear aggregation in the preceding saturation example turns out to be a general principle for all observer-invariant models. For an ecological model to be label invariant, the influence of multiple species acting through a single, shared mechanism must initially combine as a weighted sum of their abundances. This requirement applies regardless of whether populations influence each other through resource depletion, competitive stress, predation pressure or another ecological mechanism. After these initial influences are linearly summed, more complex, nonlinear ecological effects (like saturating functional responses) can be applied. This means that per capita growth rates in an observer-invariant model must take the form

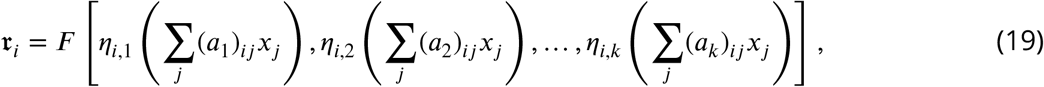

where *F* is an arbitrary function combining *k* different ecological mechanisms. Each influence, *η* _*i, m*_, is itself an arbitrary function, but its argument is critically a *linear combination* of the abundances *x*_*j*_ weighted by their specific interaction parameters (*a*_*m*_) _*ij*_ for that particular mechanism *m*. For instance, in the Lotka–Volterra model, we have *k* = 2 (intrinsic growth and competition), where *F*(*η*_*i*,1_, *η*_*i*,2_) = *η*_*i*,1_ + *η* _*i*,1_. The first pathway, intrinsic growth, is a constant function *η*_*i*,1_ (Σ _*j*_ (*a*_1_)_*ij*_*x*_*j*_) = *r*_*i*_ (as it is a property of species *i* alone); and the second pathway, competition, is an identity function *η*_*i*,2_(Σ _*j*_ (*a*_2_)_*ij*_*x*_*j*_) = Σ _*j*_ (*a*_2_)_*ij*_*x*_*j*_ (the linear sum of competitive effects).

There are two complementary ways to understand why label invariance necessitates linear building blocks. First,it emerges from the continuous space of traits (Meszéna *et al*., 2005). In this formalism, a species is represented by a Schwartz distribution—for instance, a Dirac delta function, which concentrates the entire population’s effect at a single, sharp point on the trait axis (see the mathematical details in Appendix B). The critical constraint is that the product of two such distributions is mathematically ill-defined. Therefore, to calculate the total competitive pressure on an individual, the influences of all other individuals must first be summed into a single, aggregate value. Only then can this value be used as the input for a more complex, nonlinear function that determines the final impact on per capita growth. This “sum first, then transform” logic, which ensures the model respects trait continuity, naturally leads to the structure shown in Eq. 19.

Second, a purely mathematical proof reinforces this conclusion (Ansmann & Bollenbach, 2021). It is relatively straightforward to see one side of this mathematical argument: if a model is built using linear sums, it will satisfy label invariance. The deep result established by Ansmann & Bollenbach (2021) is the converse: this linear structure is not just sufficient, but *necessary* for label invariance. In their work, the main (and mild) assumption is that the interactions are fixed and do not depend on the total number of species (regarding this last point, see Appendix G for a justification). This principle extends to more complex dynamics, such as the formulation of higher-order interactions, where seemingly equivalent mathematical forms can hide fundamentally different ecological assumptions about interaction pathways (see Appendix H for a detailed analysis).

The linear pathways constraint shows how to combine the influences of ecologically similar entities acting through a specific shared mechanism. It does not imply that every species in the entire community must appear with a nonzero coefficient in every linear sum Σ _*j*_ (*a*_*m*_)_*ij*_*x*_*j*_. Indeed, for many pairs of species (*i,k*)_*ij*_ and a given mechanism *m*, the interaction coefficient (*a*_*m*_)_*ij*_ will simply be zero, signifying that species *k* does not participate in or influence species *i* through that particular pathway. For instance, in our saturation example (Formulation 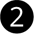), the sum *b*_1_*x*_1_ + *b*_2_*x*_2_ represents the total handling time demand imposed by *all consumed resources* on the predator’s *single processing capacity*. If the predator consumed a third resource *x*_3_ via a completely independent pathway (e.g., filter-feeding vs. seed-cracking for birds), that interaction might involve a separate linear sum relevant only to that pathway. Similarly, we would not expect the abundance of a prey species and its predator to be linearly combined to determine, for instance, the growth rate of another prey species; their ecological roles and the pathways of their influence are fundamentally different, and their respective coefficients in such a sum would reflect this by being zero.

### 4.3 Applying the principles: a label invariant multispecies model

To illustrate the linear sums principle in practice, we turn to the plant-animal mutualism model of Bastolla *et al*. (2009)—the sole model found to satisfy label invariance out of the ten surveyed by Moisset de Espanes *et al*. (2021). It describes the dynamics of interacting plant (*P*) and animal (*A*) guilds. Focusing on a representative plant species *i*, its population dynamics are given by (Bastolla *et al*., 2009):

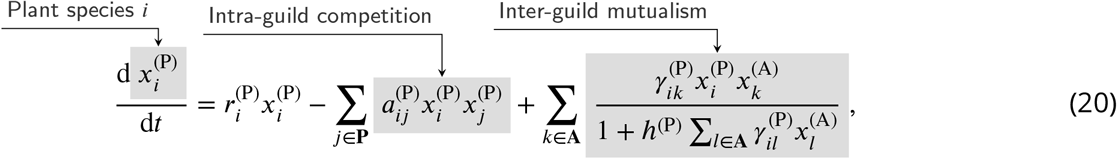

where 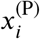 and 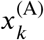 are the abundances of plant species *i* and of animal species *k*, 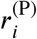 is the intrinsic growth rate of plant *i*, 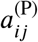 is the intra-guild competition coefficient between plant species *i* and *j*, 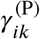 is the strength of the mutualistic interaction between plant *i* and animal *k*, and *h*^(P)^ is the handling time of the mutualistic interaction.

The formulation’s adherence to label invariance arises from two critical design choices. First, the functional forms of the intra- and inter-specific interactions are consistent with one another because of the explicit intra-guild competition term 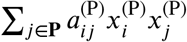. This might seem obvious, but it is a feature absent in some common mutualism models (Bascompte *et al*., 2006; Thébault & Fontaine, 2010) that might include self-limitation for a species but do not include intra-guild competition. Second, the model implements saturation effects in the inter-guild mutualism using linear aggregation. The denominator of the mutualistic term is a linear sum of the contributions from all relevant animal species, 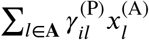 where the plant species have zero weight).

## 5 Maintaining label invariance in a noisy world

Ecologists have long grappled with incorporating stochasticity into ecological models (Lande *et al*., 2003). It is not obvious how to incorporate stochasticity into ecological models while maintaining their label invariance, especially in light of the violations we have already described. As in our discussion of how to build label invariant models in Section 4, the key insight is to consider the ecological mechanisms that generate noise, particularly how its impact scales with population size. Two principal forms of ecological stochasticity, when modeled with their mechanistic origin in mind, inherently satisfy this principle.

First, *demographic stochasticity* is the randomness inherent in the life histories of individuals— whether a particular seed germinates, an individual finds a mate, or survives an encounter (May, 1973). The magnitude of demographic fluctuations in d*x*_*i*_/d*t* scales with the square root of the population size, 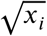. This is a direct consequence of the central limit theorem applied to summing many small, independent random events at the individual level, a scaling that also finds robust empirical support (Desharnais *et al*., 2006). Furthermore, demographic noise, arising from the unique fates of individuals, is naturally uncorrelated between distinct species. Crucially for label invariance, it is therefore uncorrelated between arbitrarily defined, identical sub-species if we are to consider their noise contributions separately. These two properties—the 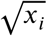 scaling of its standard deviation and its intrinsic lack of correlation across such distinct units—are what ensure that demographic stochasticity respects label invariance (see proof in Appendix I).

Second, *environmental stochasticity* captures the effects of broader-scale fluctuations—a good year for rain, a harsh winter, a widespread resource boom or bust—that tend to affect all individuals of a given type in a similar, often proportional, manner. Thus, the magnitude of environmental fluctuations in d*x*_*i*_/d_*t*_ scales directly with the population size *x*_*i*_. If we split a species *x*_0_ into identical sub-species *x*_1_ and *x*_2_, these sub-species, being ecologically indistinguishable, must experience and respond to such broad environmental shifts in precisely the same way. A widespread frost does not selectively target *x*_1_ over *x*_2_ if they are, in reality, the same. This means that the random environmental influence on their per capita growth rates must be perfectly synchronized across any such arbitrarily defined sub-groups. This requirement for perfect correlation for identical individuals stands in stark contrast to the uncorrelated nature of demographic noise. These two properties— the *x*_*i*_ -scaling of its standard deviation and the perfect correlation of the underlying noise driver for identical individuals—guarantee label invariance for environmental stochasticity.

As we prove in Appendix I, these two noise formulations are the only ones within a large class of potential noise structures that satisfy label invariance. This theoretically derived form aligns with, and provides a first-principles justification for, frameworks already widely adopted in ecological modeling (Ovaskainen & Meerson, 2010):

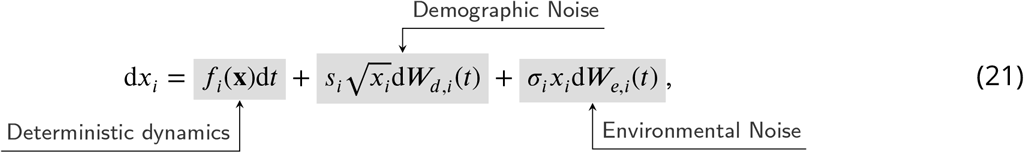

where *f*_*i*_ (x) is the observer-invariant deterministic part of species *i*’s growth, d *W*_*d,i*_ (*t*) represents the increment of a Wiener process for demographic noise with *s*_*i*_ denoting it magnitude, and d*W*_*e,i*_(*t*) represents the Wiener increment for environmental noise with *σ*_*i*_ denoting it magnitude.

Our analysis also highlights the critical, yet often overlooked, importance of the correlation structure of noise in stochastic ecological models: demographic noise must be treated as uncorrelated across arbitrarily defined units, while environmental noise must be highly correlated among similar species (this correlation is often unmodelled). The construction of more complex phenomenological stochastic models should have a clear ecological origin, for instance, by deriving them from underlying master equations of more mechanistic processes (Strang *et al*., 2019). Such derivations automatically embed the correct population scaling and the appropriate correlation properties that ensure label invariance is satisfied.

## 6 Integrating label invariance with data

Crafting models that adhere to label invariance is, as we have detailed, straightforward. Yet, theoretical coherence, however satisfying, does not guarantee empirical explanatory power. In this section, we show through a concrete example that label invariance can be used as an additional criterion in the standard model selection process when analyzing empirical data.

Annual plant are a well-suited empirical system for testing ecological theories. Ecologists have a suite of models to describe the ecological dynamics of these systems (Hart *et al*. 2018, Table 1). We consider data from field experiments in a California grassland, where fecundity and germination were measured experimentally (Van Dyke *et al*. 2022). A standard, data-first approach—employing, for instance, the Watanabe–Akaike information criterion (WAIC) for model selection—might be expected to pin down a single, best-performing model. The data, as is often the case, are more complex. As Armitage (2024) reports, three distinct models—the Beverton–Holt with an exponent, the Law–Watkinson model, and a Ricker model employing log-transformed abundances—all produce similar WAIC scores. All three appear to fit the data reasonably well.

**Table 1.** The names and functional forms of models commonly fit to annual plant data. Here, *f*_*i*_ is the fecundity of (or the number of seeds produced by) an individual of species *i. x*_*i*_ is the abundance of species *i, λ*_*i*_ is the number of seeds that species *i* produces without competition, *α*_*ij*_ measures the competitive effect of species *j* on species *i* and *β* (which is only present in the modified Beverton–Holt model) describes the functional form of the decrease in seed production with increasing competitor density. Generally, these models describe competitive systems, so *α*_*i*_ *>* 0. The WAIC (Watanabe–Akaike information criterion) are measures of model performances, where lower values correspond to better performance. These scores are adapted from Armitage (2024).

| Model ID | Model Name | Functional form | Observer Invariant? | WAIC Score |
| --- | --- | --- | --- | --- |
| 1 | Beverton-Holt with exponent | $f_i = \frac{\lambda_i}{(1+\alpha_{ii}x_i+\alpha_{ij}x_j)^\beta}$ | Yes | $382 \pm 70$ |
| 2 | Law-Watkinson | $f_i = \frac{\lambda_i}{1+x_i^{\alpha_{ii}}+x_j^{\alpha_{ij}}}$ | No | $382 \pm 70$ |
| 3 | Ricker with log abundances | $f_i = \lambda_i e^{-\alpha_{ii} \log(x_i+1) - \alpha_{ij} \log(x_j+1)}$ | No | $383 \pm 69$ |
| 4 | Beverton-Holt | $f_i = \frac{\lambda_i}{1+\alpha_{ii}x_i+\alpha_{ij}x_j}$ | Yes | $392 \pm 68$ |
| 5 | Lotka-Volterra | $f_i = \lambda_i - \alpha_{ii}x_i - \alpha_{ij}x_j$ | Yes | $423 \pm 81$ |
| 6 | Ricker | $f_i = \lambda_i e^{-\alpha_{ii}x_i - \alpha_{ij}x_j}$ | Yes | $423 \pm 8$ |

Here, label invariance provides an additional means of differentiation. Though these annual plant models are framed in discrete time, the logic of label invariance remains valid. Among the six candidates presented in Table 1, the Law–Watkinson model and the Ricker formulation employing log-transformed abundances fail the label invariance test. Their construction mirrors the structural flaws of the problematic continuous-time examples previously discussed (e.g., Model 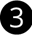 in Eq. 18).

Tellingly, this theoretical failing resonates with independent critiques from plant population biology: these same models also contravene the fundamental “law of constant yield”—a principle that emerges from the finite nature of soil resources for plant growth and seed production (Harper 1977; Van Dyke *et al*. 2024). When a model fails a test of logical consistency and contradicts established ecological regularities, its apparent statistical fit is less convincing. Importantly, the Beverton–Holt model with an exponent also does not always produce the law of constant yield, despite being label invariant, so these two criteria are not exactly equivalent.

The inferential strength of model selection is amplified when it is integrated with, and constrained by, sound theoretical principles, like label invariance or the law of constant yield. While the Law– Watkinson and the log-transformed Ricker models produce good fits, their violation of label invariance (and, in this case, other well-established ecological principles) signals potential biological inadequacy, possibly undermining their explanatory power.

## 7 Label invariance in the ecological literature

We are not the first to propose label invariance. Its core ideas, or variants thereof, have surfaced repeatedly and often independently within the ecological literature. This section aims to trace that fragmented lineage.

The Lotka–Volterra model is label invariant, but it is also one of the simplest possible descriptions of the dynamics of interacting populations. In more complex models that incorporate specific biological mechanisms, one must reckon with label invariance, leading to its independent emergence in various subfields of ecology. For example, it is a common empirical observation that predator consumption rates saturate as a function of prey density (Solomon, 1949; Holling, 1959a,b, 1965), leading ecologists to incorporate saturating functional forms into food web models (Holling, 1959b; DeAngelis *et al*., 1975; Beddington, 1975; Arditi & Akçakaya, 1990; Arditi & Ginzburg, 1989; Abrams *et al*., 2000). A natural next step was to generalize these models to food webs with multiple prey species, but there are a few different ways to do this. Early proponents of the label invariance concept showed that some of the original attempts at generalization were in fact observer variant (Arditi & Michalski, 1996; Kuang, 2002; Murrell *et al*., 2004). Conversely, adhering to label invariance guided principled derivations of food web models (Berryman *et al*., 1995; Drossel *et al*., 2001; Morozov & Petrovskii, 2013; van Leeuwen *et al*., 2013; Vallina *et al*., 2014).

Similar ideas have been formulated in the field of adaptive dynamics (Brännström *et al*., 2013). In this context, the “continuity tenet” (Waxman & Gavrilets, 2005; Meszéna, 2005) states that the relative abundance of similar phenotypes cannot have a large effect on their fitness function. The continuity tenet is a key assumption for adaptive dynamics, permitting the mathematical derivation of its central machinery. It is also relevant to the definitions of diversity metrics in conservation biology. On the face of it, one could define species diversity as simply the number of unique species, but this definition neglects the potential relatedness of different species (Weitzman, 1992; Solow *et al*., 1993; Leinster & Cobbold, 2012; Pavoine & Ricotta, 2019). The extinction of a species with no close phylogenetic relatives may be considered a greater loss than a species with many extant relatives. These considerations led previous work to propose diversity metrics that are either based on some similarity metric between species such as genetic distance (Solow *et al*., 1993), or to directly assess diversity in trait space instead of focusing on species as separate entities (Olusoji *et al*., 2023).

From evolutionary dynamics to conservation biology, researchers have grappled with how the classification of different biological types modifies the resulting understanding of the interactions between these units. Two recent works (Ansmann & Bollenbach, 2021; Moisset de Espanes *et al*., 2021) have formalized and reviewed label invariance, demonstrating how to check whether a given model satisfies it. Both approaches used sophisticated technical machinery to show that label invariant models have specific mathematical properties.

## 8 Discussion: When to worry about label invariance

Ecology has long grappled with the challenge of capturing the continuous nature of biological variation within a discrete modeling framework. Label invariance offers a principle rooted in ecological reasoning: individuals that are ecologically identical should behave identically in a model, regardless of whether they are split into distinct subpopulations or lumped into a single group. This principle emerges from trait continuity — the idea that ecological traits vary gradually in nature, and models should respect this unless there is a strong reason to violate it. Label invariance is a conceptual tool for examining model assumptions, helping to ensure that divisions between populations correspond to meaningful ecological distinctions or whether they are artifacts of model formulation or data structure.

Label invariance is not a rigid requirement that all models must adhere to, but rather a conceptual lens through which we can examine the assumptions embedded in ecological models. Models that break label invariance are not, therefore, inherently wrong, but they encode implicit ecological distinctions. For instance, in an emergent neutrality model, a failure of label invariance indicates that bill depth alone does not fully explain seed-eating bird behavior, exposing a reliance on unmodeled traits (Barabás *et al*., 2013; Ansmann & Bollenbach, 2021). Thus, this principle can serve as a diagnostic tool, revealing hidden differentiators. On the other hand, enforcing label invariance can sometimes increase model complexity. In cases where ecological entities are fundamentally distinct in their functional roles (e.g., predator and prey), encoding these distinctions through simpler model structures might be more pragmatic than formally preserving label invariance.

A productive approach is to formulate models with label invariance from the outset. This clarifies when species are considered different based on the relevant ecological coordinates, preventing the inadvertent incorporation of unrecognized structure. This practice enhances model transparency, clarifies assumptions, and ensures insights are reliably tied to ecological principles. When models preserve label invariance, emergent patterns can be confidently attributed to explicitly modeled traits and interactions, which is especially valuable for identifying general principles of coexistence, community assembly or biodiversity.

In essence, label invariance is a guiding principle rooted in trait continuity that highlights implicit assumptions and offers practical guidance for constructing clearer, more interpretable, and consistent ecological models.

## Acknowledgement

We thank Rafael D’Andrea, Kyle Dahlin and Athma Senthilnathan, and members of the Levine Lab for comments.

## Footnotes

1 For clarity, we have included additional self-regulation (*a*_11_ and *a*_22_) beyond the sublinear effects. Its presence does not affect the diversity-stability properties of this model (Hatton *et al*., 2024).

