## Supplementary Information for "Label invariance: a guiding principle for ecological models"

Table of Contents

**A Proof of label invariance for Lotka–Volterra dynamics** **S2**

**B Mathematical Formulation of Label Invariance** **S3**

**C Label invariance in consumer-resource models** **S4**

C.1 Consumer-resource model with consumption and production . . . . . . . . . . . . . S4

C.2 Consumer-resource models with saturating functional responses . . . . . . . . . . . S4

**D Proof of label invariance in ecophysiological model** **S6**

D.1 Competition for time model . . . . . . . . . . . . . . . . . . . . . . . . . . . . . S6

D.2 Root competition model . . . . . . . . . . . . . . . . . . . . . . . . . . . . . . . S7

**E Niche differences in LI and non-LI models** **S10**

E.1 Label invariant models do not have intrinsic niche differences . . . . . . . . . . . . S10

E.2 Label variance is not sufficient to have intrinsic niche differences . . . . . . . . . . S11

E.3 Niche differences analysis in the emergent neutrality model . . . . . . . . . . . . . S12

E.4 Niche differences analysis in the sublinear model . . . . . . . . . . . . . . . . . . S12

E.5 Niche differences analysis in the nonlinear feedback model . . . . . . . . . . . . . S13

E.6 Niche differences analysis in the adaptive foraging model . . . . . . . . . . . . . . S14

**F Private interaction mechanisms are necessary for a positive diversity-stability relationship: sublinear model example** **S15**

**G Scaling parameters with the number of species** **S16**

G.1 GLV with scaled interaction . . . . . . . . . . . . . . . . . . . . . . . . . . . . . S16

G.2 Adaptive foraging model . . . . . . . . . . . . . . . . . . . . . . . . . . . . . . . S16

**H Label invariance with higher-order interaction** **S18**

**I Label Invariance in Stochastic Models** **S20**

I.1 The Label Invariance Test for SDEs . . . . . . . . . . . . . . . . . . . . . . . . . S20

I.2 Proof: Testing Common Noise Formulations . . . . . . . . . . . . . . . . . . . . . S20

I.3 General Proof for Power-Law Noise . . . . . . . . . . . . . . . . . . . . . . . . . . S21

I.4 Itô vs. Stratonovich Interpretation . . . . . . . . . . . . . . . . . . . . . . . . . . S22

#### A Proof of label invariance for Lotka–Volterra dynamics

The Lotka–Volterra dynamics satisfies label invariance. Consider the Lotka–Volterra dynamics with species labeled by $i$ from 0, 3, 4, ... S:

$$\frac{\mathrm{d}x_i}{\mathrm{d}t} = r_i x_i + a_{ii} x_i^2 + \sum_{j=0;j\neq i}^{S} a_{ij} x_i x_j. \tag{S1}$$

If we split species $x_0$ into sub-species $x_1$ and $x_2$, then the dynamics governing the sub-species are

$$\frac{\mathrm{d}x_1}{\mathrm{d}t} = r_0 x_1 + a_{00} x_1^2 + \sum_{j=1}^{S} a_{ij} x_1 x_j + a_{00} x_1 x_2, \tag{S2}$$

$$\frac{\mathrm{d}x_2}{\mathrm{d}t} = r_0 x_2 + a_{00} x_2^2 + \sum_{j=1}^{S} a_{ij} x_2 x_j + a_{00} x_2 x_1, \tag{S3}$$

because the interaction between sub-species $x_1$ and $x_2$ are equal to the intraspecific interaction of species 0 (i.e., $a_{00}$). Thus, the sum of the abundances of sub-species $x_1$ and $x_2$ is

$$\underbrace{\frac{\mathrm{d}(x_1 + x_2)}{\mathrm{d}t}}_{\text{Observer 2}} = r_0(x_1 + x_2) + a_{00}(x_1 + x_2)^2 + \sum_{j=1}^{S} a_{ij}(x_1 + x_2)x_j = \underbrace{\frac{\mathrm{d}x_o}{\mathrm{d}t}}_{\text{Observer 1}}, \tag{S4}$$

which means that the GLV dynamics satisfies label invariance.

### B Mathematical Formulation of Label Invariance

Treating label invariance as a special case of trait continuity has the added benefit of immediately yielding another method for checking whether a given model adheres to these principles. Let us change perspectives and consider population size not as a function of species identity, but as a function of an individual's position in trait space. That is, instead of writing $x_i$ for the population size of a species $i$, we write a distribution $n(x)$ that is parameterized by some underlying trait(s) $x$. This means that $n(x)\,\mathrm{d}x$ is the population size of individuals with trait values falling between $x$ and $x+\mathrm{d}x$. The per capita growth rates are now also written in terms of this distribution, as $\mathfrak{r}(x, n(y))$. (The way to read this is that, for any trait $x$ and trait distribution function $n(y)$, we get back a single growth rate value. In mathematical terms, $\mathfrak{r}(x, n(y))$ is a *functional* of $n(y)$: a function that transforms a whole function into a single value.) We now require that a small $\delta n(y)$ change to $n(y)$ should induce a correspondingly small $\delta\mathfrak{r}(x, n(y))$ change in per capita growth, so that $\delta\mathfrak{r}(x, n(y))/\delta n(z) = \mathfrak{a}(x, z)$ is a finite, continuous, and well-behaved function for all trait value pairs $x$ and $z$.[2]

In particular, if there exists some trait value $x$ for which $\mathfrak{a}(x, x)$ fails to be finite, then both trait continuity and label invariance are broken: individuals with trait $x$ get preferentially (dis-)advantaged just because their trait value is what it is. As a simple example, let us take a small variant of the continuous-trait version of the Lotka–Volterra model:

$$\mathfrak{r}(x, n(y)) = r(x) - \int g(x, y) n(y)\,\mathrm{d}y - n(x). \tag{S5}$$

Here $r(x)$ is the intrinsic growth rate of individuals with trait $x$, and $g(x, y)$ is the effect of individuals (of unit population size) with trait $y$ on the per capita growth of individuals with trait $x$. Ignoring the final $-n(x)$ term, this is just a Lotka–Volterra model with $\mathfrak{a}(x, z) = g(x, z)$. Indeed,

$$\begin{aligned} \mathfrak{a}(x, z) &= \frac{\delta}{\delta n(z)}\left[r(x) - \int g(x, y) n(y)\,\mathrm{d}y\right] \\ &= -\int g(x, y)\frac{\delta n(y)}{\delta n(z)}\,\mathrm{d}y = -\int g(x, y)\delta(y - z)\,\mathrm{d}y = -g(x, z), \end{aligned} \tag{S6}$$

where $\delta(y - z)$ is the Dirac delta (a "Gaussian distribution with its variance approaching zero": it is 0 for $y \neq z$ and infinite for $y = z$, but in such a way that $\int \delta(y - z)\,\mathrm{d}y = 1$). As long as $g(x, z)$ is a well-hehaved function, the model satisfies trait continuity and label invariance. But if we also consider the $-n(x)$ term in Eq. S5, then the functional derivative becomes

$$\mathfrak{a}(x, z) = \frac{\delta\mathfrak{r}(x, n(y))}{\delta n(z)} = -g(x, z) - \frac{\delta n(x)}{\delta n(z)} = -g(x, z) - \delta(x - z). \tag{S7}$$

This is manifestly infinite for all $x = z$, breaking continuity and label invariance.

[2] A quick guide to interpreting functional derivatives like $\delta\mathfrak{r}(x, n(y))/\delta n(z)$: imagine that trait space is discrete, such that $\mathfrak{r}_i(x_j)$ is the per capita growth rate of individuals with the $i$th trait, as a function of the full trait distribution. ($x_j$ in the argument is shorthand for saying that the growth rates may depend on each of $x_1$, $x_2$, ...) If we now change the trait distribution by adding a small $\mathrm{d}x_j$ to $x_j$, we get

$$\mathfrak{r}_i(x_j + \mathrm{d}x_j) = \mathfrak{r}_i(x_j) + \sum_k \frac{\partial\mathfrak{r}_i(x_j)}{\partial x_k}\mathrm{d}x_k,$$

where the usual partial derivatives appear under the summation. The functional derivative is the generalization of this idea when the index $i$ is no longer discrete, so we use a continuous trait index $x$ instead:

$$\mathfrak{r}(x, n(y) + \delta n(y)) = \mathfrak{r}(x, n(y)) + \int \frac{\delta\mathfrak{r}(x, n(y))}{\delta n(z)}\delta n(z)\,\mathrm{d}z.$$

The quantity $\delta\mathfrak{r}(x, n(y))/\delta n(z)$ is by definition the functional derivative of $\mathfrak{r}(x, n(y))$ with respect to $n(z)$. It is the continuous analogue to the partial derivatives $\partial\mathfrak{r}_i(x_j)/\partial x_k$.

#### C Label invariance in consumer-resource models

##### C.1 Consumer-resource model with consumption and production

Consider the following model for the dynamics of $S$ consumers and $K$ resources introduced in (Butler & O'Dwyer 2018):

$$\frac{dR_i}{dt} = \rho_i - R_i \sum_{j=1}^{S} C_{ij} x_j + \sum_{j=1}^{S} P_{ij} x_j \tag{S8}$$

$$\frac{dx_i}{dt} = \epsilon x_i \sum_{j=1}^{K} C_{ji} R_j - x_i \sum_{j=1}^{K} P_{ji} - \mu_i x_i \tag{S9}$$

where $R_i$ and $x_i$ are the abundances of the $i$-th resource and consumer respectively. Resource $i$ flows abiotically into the system at rate $\rho_i$, are depleted by consumer $j$ at rate $C_{ij}$ and are also produced by consumer $j$ at rate $P_{ij}$. Consumer $i$ grows on all resources with a common efficiency $\epsilon$ and die at rate $\mu_i$. This model was originally developed for microbial communities in which resources are commonly abiotic and consumers can grow on each other's byproducts. To facilitate comparison with the other models we have considered, we employ a timescale separation for the resources (MacArthur 1970) to arrive at a set of equations for consumers alone:

$$\frac{dx_i}{dt} = \epsilon x_i \sum_{j=1}^{K} C_{ji} \frac{\rho_j + \sum_k P_{jk} x_k}{\sum_k C_{jk} x_k} - x_i \sum_{j=1}^{K} P_{ji} - \mu_i x_i \ . \tag{S10}$$

To simplify the analysis, let's consider just two consumers competing for and producing $K$ resources. Assuming that these two consumers are identical, the dynamics of consumer 1 simplify to be

$$\frac{dx_1}{dt} = \epsilon x_1 \sum_{j=1}^{K} C_{j1} \frac{\rho_j + P_{j1} x_1 + P_{j2} x_2}{C_{j1} x_1 + C_{j2} x_2} - x_1 \sum_{j=1}^{K} P_{j1} - \mu_1 x_1 \tag{S11}$$

$$= \epsilon x_1 \sum_{j=1}^{K} \frac{\rho_j + P_j (x_1 + x_2)}{x_1 + x_2} - x_1 \sum_{j=1}^{K} P_j - \mu x_1 \tag{S12}$$

where we have defined $P_j = P_{j1} = P_{j2}$ to be the common production rate between the two consumers of resource $j$ and $\mu = \mu_1 = \mu_2$. The dynamics of the sum of the two species populations is

$$\frac{d(x_1 + x_2)}{dt} = \epsilon (x_1 + x_2) \sum_{j=1}^{K} \frac{\rho_j + P_j (x_1 + x_2)}{x_1 + x_2} - (x_1 + x_2) \sum_{j=1}^{K} P_j \mu (x_1 + x_2) = \frac{dx_0}{dt} \tag{S13}$$

which is the same as the dynamics for the joint species $x_0$, meaning this model is label invariant. This argument relies on the additivity of the specific functional forms we used to represent consumption and production. For example, if production depended on consumer abundances with a type II saturating functional response, then we would not be able to group the consumer abundances together in equation (S12) as we have done, violating label invariance.

##### C.2 Consumer-resource models with saturating functional responses

In foundational work, Monod (1949) observed that bacterial consumption of a given nutrient saturated with the density of that nutrient. In generalizing this behavior to the consumption of multiple resources, ecologists have often simply summed these Monod growth forms together for each resource (León & Tumpson 1975). This intuitive generalization violates label invariance in a similar way that the models we presented in the main text with a saturating functional response did. Consider a simple model of one consumer consuming one resource:

$$\frac{dR}{dt} = R(r - aR) - f(R)CN \tag{S14}$$

$$\frac{dN}{dt} = \epsilon NCf(R) - \mu N \tag{S15}$$

where $R$ is the abundance of the resource, $N$ is the abundance of the consumer, $r$ is the intrinsic growth rate of the resource, $a$ is the strength of intraspecific competition for the resource, $C$ is the consumption of the resource by the consumer, $f$ is the functional form of resource consumption and $\epsilon$ and $\mu$ are analogous to the previous section. Splitting the resources into two identical populations, the dynamics of their sum is

$$\frac{d(R_1 + R_2)}{dt} = r(R_1 + R_2) - a(R_1 + R_2)^2 - CN(f(R_1) + f(R_2)) \tag{S16}$$

so that if $f(R_1) + f(R_2) = f(R_1 + R_2)$ then this model is label invariant. If each of the consumers follow a standard saturating functional response, $f(R_1) = \frac{R}{K+R_1}$, then this equality is not satisfied and the model is observer variant. If instead the denominator has both resources (ie. $f(R_1) = \frac{R}{K+R_1+R_2}$, then model is label invariant. This result has a clear biological interpretation. In a microbial example, similar resources are processed through the same internal metabolic pathways, so vast amounts of any of the resources depleted via such a pathway will decrease the rate of consumption of all resources. This calculation also suggests that more similar resources should not only be consumed at similar rates but also through more similar functional forms. More careful consideration of the biochemistry of resource consumption should naturally give rise to such similarity.

### D Proof of label invariance in ecophysiological model

#### D.1 Competition for time model

Here we prove that the ecophysiological model for competition among annual plants developed by Levine *et al.* (2022) is label invariant. The model is formulated in discrete time, and its core competitive mechanism is the collective drawdown of a shared water pool, which shortens the growing season for all species.

The population dynamics for a species $i$ from year $T$ to $T+1$ are given by Eq. S14 in the supplement:

$$\frac{N_{i,T+1}}{N_{i,T}} = FG_i^{L+1}\tau_i(\mathbf{N}_T) \tag{S17}$$

where $N_{i,T}$ is the population density, $F$ is the fecundity constant, $G_i^{L+1}$ is the species-specific biomass growth rate, and $\tau_i(\mathbf{N}_T)$ is the actual growing season length on a transformed timescale. The proof of label invariance hinges on demonstrating that $\tau_i$ for every species is unaffected by the splitting of another species into identical sub-species.

The growing season length $\tau_i$ is the time it takes for the soil water content to drop to species $i$'s critical threshold $W_i^*$. The dynamics of water consumption are central. For the first species to finish its growing season (species 1, with the highest $W_1^*$), its season length $\tau_1$ is given by Eq. S12 in the supplement:

$$\tau_1 = \left(\frac{L+1}{E}\right)\frac{W_0 - W_1^*}{\sum_{j=1}^{Q} N_{j,T}G_j^L} \tag{S18}$$

where $W_0$ is initial water content, $E$ is a transpiration constant, $L$ is an allometric exponent, and the denominator represents the total water consumption pressure from the entire community. For any subsequent species $i$, its growing season is determined piecewise, but the rate of water drawdown in any given time interval is always proportional to a linear sum of the densities of the species that are still actively growing. The denominator in Eq. S18 is the key to the proof.

We now perform the label invariance test step-by-step. First, consider an initial state with a community of $Q$ species with densities $\mathbf{N} = \{N_1, \ldots, N_Q\}$. The total water consumption pressure term in the denominator of Eq. S18 is:

$$C_{total} = \sum_{j=1}^{Q} N_{j,T}G_j^L \tag{S19}$$

Second, we split one species, $k$, into two identical sub-species, $k_1$ and $k_2$. The new community formally has $Q+1$ species. For the split to be valid, the sub-species must be identical to the original, which imposes the following conditions:

- Density is conserved: $N_k = N_{k,1} + N_{k,2}$.
- Traits are identical: $G_k^L = G_{k,1}^L = G_{k,2}^L$ and $G_k^{L+1} = G_{k,1}^{L+1} = G_{k,2}^{L+1}$.
- Critical thresholds are identical: $W_k^* = W_{k,1}^* = W_{k,2}^*$ and $\tau_k^* = \tau_{k,1}^* = \tau_{k,2}^*$.

Third, we calculate the new total water consumption pressure, $C'_{total}$, for the community of $Q+1$ species. We separate the sum into the species that were not split and the new sub-species:

$$C'_{total} = \left(\sum_{j=1, j\neq k}^{Q} N_j G_j^L\right) + N_{k,1}G_{k,1}^L + N_{k,2}G_{k,2}^L$$

Applying the conditions for an identical split, this becomes:

$$\begin{aligned}
C'_{total} &= \left( \sum_{j=1, j\neq k}^{Q} N_j G_j^L \right) + N_{k,1} G_k^L + N_{k,2} G_k^L \\
&= \left( \sum_{j=1, j\neq k}^{Q} N_j G_j^L \right) + (N_{k,1} + N_{k,2}) G_k^L \\
&= \left( \sum_{j=1, j\neq k}^{Q} N_j G_j^L \right) + N_k G_k^L \\
&= \sum_{j=1}^{Q} N_j G_j^L = C_{total}
\end{aligned} \tag{S20}$$

The total water consumption pressure term is mathematically identical before and after the split. Since this term is the only part of the equation for $\tau_i$ that depends on community composition, the growing season length $\tau_i$ for all species remains unchanged.

Consequently, the dynamics of all other species ($j \neq k$) are unaffected. The total population of the split species $k$ also follows the original dynamics. Since their per capita growth rates are identical, the total population in the next year is:

$$\begin{aligned}
N_{k,T+1} &= N_{k_1,T+1} + N_{k_2,T+1} \\
&= N_{k_1,T}\left(F G_k^{L+1} \tau_k\right) + N_{k_2,T}\left(F G_k^{L+1} \tau_k\right) \\
&= (N_{k_1,T} + N_{k_2,T})\left(F G_k^{L+1} \tau_k\right)
\end{aligned} \tag{S21}$$

This is identical to the population dynamics of the unsplit species $k$. The model is therefore label invariant because the competitive impact on the shared water resource is determined by a linear summation of species densities, weighted by their traits. This property is also evident in the decomposition of per capita competitive effects (Supplement Section 3.6), where the derived intra- and inter-specific competition coefficients ($\alpha_{ii}$ and $\alpha_{ij}$) are shown to be constructed from the same fundamental growth parameters, ensuring that the interaction between identical sub-species is equivalent to their self-interaction.

#### D.2 Root competition model

Here we prove that the consumer-resource model for exploitative root competition developed by Cabal *et al.* (2020) is label invariant. The model's core mechanism is that plants compete indirectly by depleting a shared, spatially explicit resource pool (e.g., water). The competitive effect of any root on the resource pool is determined solely by its density and physiological uptake rate, not by the identity of the plant it belongs to. We will show that this structure, which relies on a linear summation of effects, inherently satisfies the principle of label invariance.

The model assumes that at any given soil coordinate $\ell$, the equilibrium resource concentration $W(\ell)$ is determined by a constant input rate $I$, a physical loss rate $\delta$, and the total uptake by the roots of all $n$ plants present. From Eq. 1.1.4 in the supplementary material, this is:

$$W(\ell) = \frac{I}{\delta + \sum_{i=1}^{n} \alpha_i R_i(\ell)} \tag{S22}$$

where $R_i(\ell)$ is the root density of plant $i$ at coordinate $\ell$, and $\alpha_i$ is its per capita resource uptake rate.

The fitness of a plant $j$ at coordinate $\ell$, $G_j(\ell)$, is determined by the balance between the benefit from resource uptake and the cost of root production and maintenance, $C_j(\ell)$. From Eq. 1.1.5 in the supplement:

$$G_j(\ell) = \alpha_j R_j(\ell) W(\ell) - C_j(\ell) R_j(\ell) = \frac{\alpha_j R_j(\ell) I}{\delta + \sum_{i=1}^{n} \alpha_i R_i(\ell)} - C_j(\ell) R_j(\ell) \tag{S23}$$

The proof of label invariance rests on demonstrating that the total resource uptake term in the denominator, $\sum_{i=1}^{n} \alpha_i R_i(\ell)$, remains unchanged when a single plant is arbitrarily divided into identical sub-plants.

We now perform the label invariance test step-by-step:

1. **Initial State:** Consider a community of $n$ plants. The total resource uptake pressure at coordinate $\ell$ is given by the sum in the denominator of Eq. S22:

$$U_{total}(\ell) = \sum_{i=1}^{n} \alpha_i R_i(\ell)$$

2. **The Split:** We take one plant, $k$, and arbitrarily divide it into two identical sub-plants, $k_1$ and $k_2$. The new community now formally contains $n+1$ plants. For the sub-plants to be identical to the original, the split must satisfy the following conditions:
   - Root density is conserved at every location: $R_k(\ell) = R_{k,1}(\ell) + R_{k,2}(\ell)$.
   - Physiological traits are identical: $\alpha_k = \alpha_{k,1} = \alpha_{k,2}$.
   - The cost functions are identical: $C_k(\ell) = C_{k,1}(\ell) = C_{k,2}(\ell)$.

3. **Calculate New Total Uptake Pressure:** We compute the total resource uptake pressure, $U'_{total}(\ell)$, for the new $n+1$ plant community by separating the sum into the non-split plants and the two new sub-plants:

$$U'_{total}(\ell) = \left( \sum_{i=1, i\neq k}^{n} \alpha_i R_i(\ell) \right) + \alpha_{k,1} R_{k,1}(\ell) + \alpha_{k,2} R_{k,2}(\ell)$$

4. **Apply Split Conditions:** We substitute the identities from the split into the equation:

$$\begin{aligned} U'_{total}(\ell) &= \left( \sum_{i=1, i\neq k}^{n} \alpha_i R_i(\ell) \right) + \alpha_k R_{k,1}(\ell) + \alpha_k R_{k,2}(\ell) \\ &= \left( \sum_{i=1, i\neq k}^{n} \alpha_i R_i(\ell) \right) + \alpha_k (R_{k,1}(\ell) + R_{k,2}(\ell)) \\ &= \left( \sum_{i=1, i\neq k}^{n} \alpha_i R_i(\ell) \right) + \alpha_k R_k(\ell) \\ &= \sum_{i=1}^{n} \alpha_i R_i(\ell) = U_{total}(\ell) \end{aligned}$$

Since $U'_{total}(\ell) = U_{total}(\ell)$, the denominator in the resource equation (Eq. S22) is unchanged. This means the equilibrium resource concentration $W(\ell)$ is identical before and after the split. Consequently, for any other plant $j \neq k$, its fitness-generating function $G_j(\ell)$ (from Eq. S23) remains the same, and its optimal root allocation strategy $R_j^*(\ell)$ will be unaffected.

For the split plant itself, the sub-plants $k_1$ and $k_2$ will each optimize their own fitness. In the ESS framework, this means they will satisfy $\partial G_{k,1}/\partial R_{k,1} = 0$ and $\partial G_{k,2}/\partial R_{k,2} = 0$. Since they are identical and face an identical resource environment $W(\ell)$, their combined optimal root density $R^*_{k,1}(\ell) + R^*_{k,2}(\ell)$ will be the same as the original optimal density $R^*_k(\ell)$.

In conclusion, the model is label invariant because competitive interactions are mediated solely through the linear, additive depletion of a shared resource. The identity of a root's owner is irrelevant to its effect on the resource pool; only its density and uptake rate matter. This structure inherently satisfies the conditions for label invariance.

### E Niche differences in LI and non-LI models

Let us introduce the definition of niche difference introduced in (Spaak & De Laender 2020). Consider the case of two interacting species, $i$ and $j$:

$$\frac{1}{x_i}\frac{\mathrm{d}x_i}{\mathrm{d}t} = f_i(x_i, x_j), \tag{S24}$$

where $f_i(x_i, x_j)$ is the per capita growth rate of species $i$, and we have the equivalent equation for species $j$. The definition of niche difference for the two species is given by

$$\mathcal{N}_i \equiv \frac{f_i(0, x_j^*) - f_i(c_j x_j^*, 0)}{f_i(0, 0) - f_i(c_j x_j^*, 0)}, \tag{S25}$$

where $x_j^*$ indicates the equilibrium value of species $j$ in absence of species $i$, and the parameter $c_j$ convert the density of species $j$ in the equivalent of species $i$ needed to obtain the same effect (Spaak & De Laender 2020). To compute $c_j$ (and $c_i$) we can use the symmetry of the niche difference

$$\mathcal{N}_i = \mathcal{N}_j, \tag{S26}$$

and the fact that converting back to the original species should result in the original density

$$c_i c_j = 1. \tag{S27}$$

As an example, for Lotka–Volterra we have

$$f_i(x_i, x_j) = 1 - (a_{ii}x_i + a_{ij}x_j), \tag{S28}$$

therefore the niche difference is

$$\mathcal{N}_i = 1 - \frac{a_{ij}x_j^*}{a_{ii}c_j x_j^*} = 1 - \frac{a_{ij}}{a_{ii}c_j}. \tag{S29}$$

the conversion factor can be computed using Eqs. (S26) and (S27) giving

$$c_j = \sqrt{\frac{a_{ij}a_{jj}}{a_{ji}a_{ii}}}, \tag{S30}$$

and therefore

$$\mathcal{N}_i = 1 - \sqrt{\frac{a_{ij}a_{ji}}{a_{ii}a_{jj}}}. \tag{S31}$$

#### E.1 Label invariant models do not have intrinsic niche differences

In this section we show that in label invariant models the niche difference between two identical populations, i.e., populations with identical traits, is always 0.

An label invariant model can be written in general as

$$\frac{1}{x_i}\frac{\mathrm{d}x_i}{\mathrm{d}t} = F\left[\eta_{i,1}\left(\sum_j (a_1)_{ij}x_j\right), \eta_{i,2}\left(\sum_j (a_2)_{ij}x_j\right), \ldots, \eta_{i,k}\left(\sum_j (a_k)_{ij}x_j\right)\right], \tag{S32}$$

We first reduce to the case in which the per capita growth rate is characterized by a single process

$$\frac{1}{x_i}\frac{\mathrm{d}x_i}{\mathrm{d}t} = \eta_i\left(\sum_j a_{ij}x_j\right), \tag{S33}$$

and therefore, for two species we have

$$\frac{1}{x_i}\frac{\mathrm{d}x_i}{\mathrm{d}t} = \eta_i(a_{ii}x_i + a_{ij}x_j). \tag{S34}$$

This structure of the per capita growth rates implies that

$$\frac{1}{x_i}\frac{\mathrm{d}x_i}{\mathrm{d}t} = f_i(x_i, x_j) = f_i\left(x_i + \frac{a_{ij}}{a_{ii}}x_j, 0\right) = f_i\left(0, \frac{a_{ii}}{a_{ij}}x_i + x_j\right). \tag{S35}$$

Therefore, in general, we have explicitly

$$\mathcal{N}_i = \frac{f_i\Big((a_{ij}/a_{ii})x_j^*, 0\Big) - f_i(c_jx_j^*, 0)}{f_i(0,0) - f_i(c_jx_j*, 0)}. \tag{S36}$$

If two species have the same traits, $a_{ij}/a_{ii} = 1$ and $c_j = 1$, and therefore $\mathcal{N}_i = 0$.

In general, we have, for two species

$$\frac{1}{x_i}\frac{\mathrm{d}x_i}{\mathrm{d}t} = F\{\eta_{i,1}[(a_1)_{ii}x_i + (a_1)_{ij}x_j)], \ldots, \eta_{i,k}[(a_k)_{ii}x_i + (a_k)_{ij}x_j]\}. \tag{S37}$$

In this case, we cannot write the equivalent of Eq. (S35), because there are multiple ecological processes, each characterized by different parameters $a$. However, in full generality, we can write

$$\frac{1}{x_i}\frac{\mathrm{d}x_i}{\mathrm{d}t} = F\left\{\eta_{i,1}\left[(a_1)_{ii}\left(x_i + \frac{(a_1)_{ij}}{(a_1)_{ii}}x_j\right) + 0\right], \ldots, \eta_{i,k}\left[(a_k)_{ii}\left(x_i + \frac{(a_k)_{ij}}{(a_k)_{ii}}x_j\right) + 0\right]\right\}. \tag{S38}$$

If the two sub-population are the same species, we must have $(a_1)_{ij}/(a_1)_{ii} = (a_2)_{ij}/(a_2)_{ii} = \cdots = (a_k)_{ij}/(a_k)_{ii} = 1$, and therefore

$$f_i(x_i, x_j) = f_i(x_i + x_j, 0),$$

leading to $\mathcal{N}_i = 0$. This intuitive general result stems from the fact that in label invariant models, every ecological trait that can distinguish two different species is explicitly parametrized, and densities always appear in linear combinations.

#### E.2 Label variance is not sufficient to have intrinsic niche differences

In this section, we show, by means of a counterexample, that the violation of label invariance in a model is not sufficient to imply the existence of hidden niches in the model.

Let us consider the modified Lotka-Volterra model in which the dependence on the density of the other species is nonlinear

$$f_i(x_i, x_j) = 1 - (a_{ii}x_i^\theta + a_{ij}x_j^\theta), \tag{S39}$$

and

$$f_j(x_j, x_i) = 1 - (a_{jj}x_j^\theta + a_{ji}x_i^\theta), \tag{S40}$$

where the strength of the nonlinearity $\theta \neq 1$ is the same for both species. This model clearly violates label invariance; however, the niche difference

$$\mathcal{N}_i = 1 - \frac{a_{ij}(x_j^*)^\theta}{a_{ii}(c_jx_j^*)^\theta} = 1 - \frac{a_{ij}}{a_{ii}c_j^\theta}, \tag{S41}$$

vanishes if the two populations are the same species, i.e., $a_{ij} = a_{ii}$ and $c_j = 1$.

##### E.3 Niche differences analysis in the emergent neutrality model

The emergent neutrality model for two species can be written as

$$\frac{1}{x_i}\frac{\mathrm{d}x_i}{\mathrm{d}t} = r_i - \frac{r_i}{K_i}\left(x_i + x_j \exp\left[-\frac{w_{ij}}{\sigma^2}\right]\right) - \left(\frac{g}{x_i^2 + H^2}\right)x_i. \tag{S42}$$

The niche difference is

$$\mathcal{N}_i = 1 - \frac{\frac{r_i}{K_i}x_j^* \exp\left[-\frac{w_{ij}}{\sigma^2}\right]}{\left(\frac{r_i}{K_i} + \frac{g}{(c_j x_j^*)^2 + H^2}\right)c_j x_j^*}. \tag{S43}$$

If the two species are the same, we have $w_{ij} = 0$, $c_i = c_j = 1$, $r_i = r_j = r$ and $K_i = K_j = K$, leading to

$$\mathcal{N}_i = 1 - \frac{1}{1 + \frac{K}{r}\frac{g}{(x^*)^2 + H^2}}, \tag{S44}$$

which demonstrates irreducible niche differentiation. The niche difference vanishes only in the absence of the extra self-regulation term, i.e., for $g = 0$ or $H \to \infty$.

##### E.4 Niche differences analysis in the sublinear model

In this section, we analyze niche differences in the sublinear model and in analogous models with similar diversity-stability properties.

###### E.4.1 Sublinear model

Let us consider a variation of the sublinear model in the main text that prevents diverging invasion growth rate but retains the diversity-stability properties (Hatton *et al.* 2024)

$$\frac{1}{x_i}\frac{\mathrm{d}x_i}{\mathrm{d}t} = \Theta(1 - x_i) + \Theta(x_i - 1)x_i^{k-1} - z_i - a_{ij}x_j, \tag{S45}$$

where $k < 1$, $\Theta(x)$ is the Heaviside theta step function, and we can assume for simplicity $z_i = z_j < 1$. In this case, we have that the equilibrium in the absence of the other species is

$$x_i^* = x_j^* = z^{1/(k-1)} > 1. \tag{S46}$$

The niche difference is

$$\mathcal{N}_i = 1 - \frac{a_{ij}x_j^*}{1 - (c_j x_j^*)^{k-1}}, \tag{S47}$$

and we immediately realize that even in the case in which the two populations share the same traits, $a_{ij} = a_{ji} = a$ and $c_j = 1$, there is still a residual niche difference

$$\mathcal{N}_i = 1 - \frac{a z^{1/(k-1)}}{1 - z}. \tag{S48}$$

###### E.4.2 Theta-logistic model

It is instructive to consider a generalization of this model, i.e., a model with theta-logistic intraspecific interaction and logistic interspecific interactions

$$\frac{1}{x_i}\frac{\mathrm{d}x_i}{\mathrm{d}t} = 1 - a_{ii}x_i^{\theta} - a_{ij}x_j, \tag{S49}$$

with the same $\theta$ for both populations. For $\theta < 1$ this model enjoys the same diversity-stability properties of the sublinear model (Hatton *et al.* 2024).

The niche difference for the two species in this model is

$$\mathcal{N}_i = 1 - \frac{a_{ij}(x_j^*)^{1-\theta}}{a_{ii}c_j^\theta}. \tag{S50}$$

After computing the conversion factor

$$c_j = \left(\frac{a_{jj}a_{ij}(x_j^*)^{1-\theta}}{a_{ii}a_{ji}(x_i^*)^{1-\theta}}\right)^{1/(2\theta)}, \tag{S51}$$

we can write

$$\mathcal{N}_i = 1 - \sqrt{\frac{a_{ij}a_{ji}}{a_{ii}a_{ji}}}(x_j^*x_i^*)^{(1-\theta)/2}. \tag{S52}$$

The expression for the niche difference reduces correctly to the Lotka-Volterra case for $\theta = 1$. For $\theta \neq 1$, even in the case of identical subpopulations, $a_{ij} = a_{ii} = a_{ji} = a_{jj} = a$ we have an irreducible niche difference

$$\mathcal{N}_i = 1 - a^{(\theta-1)/\theta}. \tag{S53}$$

Considering the case $\theta = 1/2$ for analytical tractability, we also show that the niche difference between any pair of species in a community of $S$ identical species is well approximated by

$$\mathcal{N}_i \approx 1 - [a(S-2)]^{-1}, \tag{S54}$$

for $S \gg 1$ and $0 < a < 1$. Therefore, the niche overlap systematically decreases (as $1/S$) with an increasing number of species $S$.

##### E.5 Niche differences analysis in the nonlinear feedback model

The nonlinear feedback model for two species is

$$\frac{1}{x_i}\frac{\mathrm{d}x_i}{\mathrm{d}t} = r_i[K_i - x_i - g(a_{ii}x_i + a_{ij}x_j)], \tag{S55}$$

where $g(u)$ is a non-decreasing function of its argument. The authors show that when $g(u)$ is a saturating function, e.g., a Hill function of the form

$$g(u) = \frac{2\alpha u}{\alpha + 2|u|}, \tag{S56}$$

where the parameter $\alpha$ sets the saturation, it stabilizes communities, leading to more diversity than in the absence of saturation with respect to the equivalent generalized Lotka-Volterra model.

The niche difference between the two species reads in the general formulation

$$\mathcal{N}_i = 1 + \frac{g(a_{ij}x_j^*)}{c_jx_j^* - g(c_ja_{ii}x_j^*)}. \tag{S57}$$

If we choose, as a proof of concept, the Hill function in Eq (S56) and consider the case of identical species, i.e., $c_i = c_j = 1$ and $a_{ij} = a \ \forall i, j$, we find

$$\mathcal{N}_i = 1 + \frac{1}{\frac{\alpha+2|ax^*|}{2\alpha a} - 1}, \tag{S58}$$

which cannot vanish for any value of $x^*$, implying an irreducible niche differentiation.

Notice that the problem is not in the details of the nonlinearity but in the linear feedback present only for the focal species.

##### E.6 Niche differences analysis in the adaptive foraging model

We first consider the case in which the trophic link between two species is mutually exclusive, i.e., every species can either be a consumer or a resource for another one (Kondoh 2003),

$$\frac{dx_i}{dt} = x_i \left( r_i - s_i x_i + \sum_{j \neq i} (e_{ij} f_{ij} a_{ij} - f_{ji} a_{ji}) x_j \right), \tag{S59}$$

if we assume without loss of generality $f_{ij} = e_{ij} = 1 \ \forall i, j$ we have

$$\frac{dx_i}{dt} = x_i \left( r_i - s_i x_i + \sum_{j \neq i} \alpha_{ij} x_j \right), \tag{S60}$$

with

$$\alpha_{ij} = a_{ij} - a_{ji}, \tag{S61}$$

antisymmetric. Therefore, if two species are the same, we can set $\alpha_{ij} = 0$, but then the two species are non-interacting.

In the case in which loops and cannibalism are possible (Brose *et al.* 2003), we can consider the case of two species, $i$ and $j$, both predating on themselves and on each other

$$\frac{1}{x_i} \frac{\mathrm{d}x_i}{\mathrm{d}t} = 1 - a_{ii} x_i - a_{ij} x_j, \tag{S62}$$

$$\frac{1}{x_j} \frac{\mathrm{d}x_j}{\mathrm{d}t} = 1 - a_{jj} x_j - a_{ji} x_i, \tag{S63}$$

where we set $r_i = r_j = 1$ and $s_i = s_j = 0$ for simplicity, with the constraints

$$a_{ii} + a_{ij} = a_{jj} + a_{ji} = 1. \tag{S64}$$

We therefore have a Lotka-Volterra model, with niche difference

$$\mathcal{N}_i = 1 - \sqrt{\frac{a_{ij} a_{ji}}{a_{ii} a_{jj}}}, \tag{S65}$$

If the two species are the same, then $a_{ii} = a_{ij} = a_{jj} = a_{ji} = a$, resulting in vanishing niche differentiation. However, the constraints should also be satisfied at all times, giving $a = 1/2$ regardless of the population of the two species, or resource profitability (Kondoh 2003), which depends on the populations. In other words, to have adaptive foraging, two identical sub-populations should admit an irreducible niche differentiation.

### F Private interaction mechanisms are necessary for a positive diversity-stability relationship: sublinear model example

Consider the following label invariant version of the sublinear model

$$\frac{\mathrm{d}x_i}{\mathrm{d}t} = x_i \left[ \left( \sum_{j=1}^{S} b_{ij} x_j \right)^{k-1} - \sum_{j=1}^{S} a_{ij} x_j \right], \tag{S66}$$

and let us focus on the case of homogeneous interactions, with $a_{ii} = b_{ii} = 1$, and $a_{ij} = \mu_a$, $b_{ij} = \mu_b \; \forall j \neq i$.

The equilibrium abundance value, common across species, is

$$x = \left[ \frac{1 + \mu_a}{(1 + \mu_b)^{k-1}} \right]^{1/(k-2)}. \tag{S67}$$

The diagonal and off-diagonal components of the Jacobian at equilibrium are, respectively

$$J_{ii}^* = x \Big[ (k-1)(1 + \mu_b (S-1) x)^{k-2} - 1 \Big], \tag{S68}$$

and

$$J_{ij} = x \Big[ (k-1)(1 + \mu_b (S-1) x)^{k-2} \mu_b - \mu_a \Big], \tag{S69}$$

The largest eigenvalue is given by $\lambda = J_{ii}^* - J_{ij}^*$, therefore the stability condition is

$$(k-1)(1 - \mu_b + \mu_b (S-1) x)^{k-2} - (1 - \mu_a) < 0, \tag{S70}$$

which, by using the equilibrium solution, reads

$$(k-1)(1 - \mu_b) \frac{1 + \mu_a (S-1)}{1 + \mu_b (S-1)} < 1 - \mu_a, \tag{S71}$$

which becomes independent of $S$, at large $S$, as for GLV. We can also see that we need at least one of $\mu_a$ or $\mu_b$ to be $< 1$ in order for the system to be stable. Only if we choose the singular condition $\mu_b = 0$ do we obtain a positive diversity-stability relationship.

### G Scaling parameters with the number of species

A particularly subtle case of observer variance concerns models apparently built using linear combinations of densities as building blocks but where the parameters depend in some way on the number of species in the model. This dependence introduces nonlinearities, breaking label invariance. This violation can be important when analyzing how models with scaled parameters behave as a function of the number of species, but they are less relevant with a species pool of fixed size.

#### G.1 GLV with scaled interaction

Consider the GLV model for $S$ species with random interactions scaling in such a way that the effective competitive pressure felt by each species from the rest of the pool is independent of $S$

$$\frac{\mathrm{d}x_i}{\mathrm{d}t} = x_i\left(r_i - \sum_{j=1}^{S} a_{ij}x_j\right) = x_i\left(r_i - \frac{\mu}{S}\sum_{j=1}^{S} x_j - \frac{\sigma}{\sqrt{S}}\sum_{j=1}^{S}\alpha_{ij}x_j\right), \tag{S72}$$

with $\alpha_{ij} \sim \mathcal{N}(0, 1)$. This model is not label invariant because the dependency on the number of species in the parameters introduces a nonlinearity, which can be shown to be in contrast with the rule of using only a linear combination of species as building blocks for the dynamical equations. Indeed,

$$S \equiv \lim_{k\to 0}\sum_{j=1}^{S} x_j^k \equiv \sum_{j=1}^{S} x_j^0. \tag{S73}$$

is a simple nonlinear function that counts the number of species in the community, regardless of whether they have been excluded.

To show the observer variance of this model more concretely, consider the simple, neutral case of $\sigma = 0$ and $r_i = r$ for every $i$, and let us start with one species, $S = 1$,

$$\frac{\mathrm{d}x}{\mathrm{d}t} = x(r - \mu x), \tag{S74}$$

which, scaling the interaction as $\mu/S$, becomes, if we split in two species $x \equiv x_1 + x_2$

$$\frac{\mathrm{d}x_1}{\mathrm{d}t} = x_1\left[r - \frac{\mu}{2}x_1 - \frac{\mu}{2}x_2\right], \tag{S75}$$

$$\frac{\mathrm{d}x_2}{\mathrm{d}t} = x_2\left[r - \frac{\mu}{2}x_2 - \frac{\mu}{2}x_1\right], \tag{S76}$$

$$\tag{S77}$$

and, therefore, we obtain

$$\frac{\mathrm{d}(x_1 + x_2)}{\mathrm{d}t} = (x_1 + x_2)\left[r - \frac{\mu}{2}(x_1 + x_2)\right] \neq x(r - \mu x). \tag{S78}$$

#### G.2 Adaptive foraging model

The same argument is behind the lack of label invariance of the adaptive foraging model

$$\frac{\mathrm{d}x_i}{\mathrm{d}t} = x_i\left(r_i - s_i x_i + \sum_{j\in \text{ resources}} e_{ij} f_{ij} a_{ij} x_j - \sum_{j\in \text{ consumers}} f_{ji} a_{ji} x_j\right), \tag{S79}$$

with the constraint

$$\sum_{j \in \text{ resources}} a_{ij} = 1. \tag{S80}$$

Indeed, we can write

$$a_{ij} \equiv \frac{b_{ij}}{\sum_j b_{ij}} \equiv \frac{b_{ij}}{\sum_j b_{ij} x_j^0}, \tag{S81}$$

where the $b_{ij}$ are not constrained.

### H Label invariance with higher-order interaction

We then turn our attention to the higher-order interactions, where a third species modifies the pairwise interaction between two others (Kleinhesselink *et al.* 2022). While adding 3-way interaction terms to the Lotka–Volterra model might seem intuitive, seemingly identical mathematical forms can hide key differences in ecological interpretation. Consider a 3-species community ($x_1$, $x_2$, $x_3$) and focus on the dynamics of species $x_1$ in two scenarios. In the first scenario, the way species $x_3$ modifies the interaction between $x_1$ and $x_2$ differs from how $x_2$ modifies the interaction between $x_1$ and $x_3$ (Bairey *et al.* 2016). The Lotka–Volterra dynamics with higher-order interactions are written as:

$$\frac{\mathrm{d}x_1}{\mathrm{d}t} = r_1 x_1 - \overbrace{a_{11}x_1^2}^{\text{Intraspecific}} - \underbrace{a_{12}x_1x_2 - a_{13}x_1x_3}_{\text{Interspecific}} - \overbrace{\sum_{\boldsymbol{i=1}}^{\mathbf{3}}\sum_{\boldsymbol{j=1}}^{\mathbf{3}} a_{1ij}x_1x_ix_j}^{\text{Higher-order interaction}} . \tag{S82}$$

where the higher-order interaction term $a_{1ij}$ represents how presence of species $j$ affects the pairwise interaction between species 1 and $i$. We include both $a_{1\mathbf{23}}x_1x_2x_3$ and $a_{1\mathbf{32}}x_1x_3x_2$ to reflect the distinct modification paths.

In the second scenario, we do not distinguish the two paths of interaction modification and consider them as one path (Letten & Stouffer 2019). This leads to:

$$\frac{\mathrm{d}x_1}{\mathrm{d}t} = r_1 x_1 - \overbrace{a_{11}x_1^2}^{\text{Intraspecific}} - \underbrace{a_{12}x_1x_2 - a_{13}x_1x_3}_{\text{Interspecific}} - \overbrace{\sum_{\boldsymbol{i=1}}^{\mathbf{3}}\sum_{\boldsymbol{j\geq i}} a_{1ij}x_1x_ix_j}^{\text{Higher-order interaction}} \tag{S83}$$

where the higher-order interaction term $a_{1ij}$ represents the joint effect of species $i$ and $j$ on the growth of species 1. Note that $a_{132}x_1x_3x_2$ is absent in the equation, as they are counted in the term $a_{123}x_1x_2x_3$.

Why should we care about these differences in model formulation, given they are mathematically equivalent (by summing $(a_{1ij} + a_{1ji})$ in Eqn. S82 to get the new $a_{1ij}$ in Eqn. S83). However, these formulations embody fundamentally different ecological assumptions with the potential for inconsistency. Label invariance helps us expose this inconsistency. We found that only the first formulation (Eqn. S82) fully satisfies label invariance, indicating a constraint that the second formulation lacks (Eqn. S83). The presence of higher-order intraspecific interaction term $(a_{111}x_1^3)$ is the key issue. Heuristically, this cubic self-regulation can only arise from the multiple interaction modification pathways between species, which the second formulation implicitly disallows. For a detailed proof, consider three species $x$, $y$, and $z$,

$$\begin{aligned}\frac{\mathrm{d}x}{\mathrm{d}t} =& r_x x - a_{xx}x^2 - a_{xxx}x^3 - a_{xy}xy - a_{xz}xz && \text{(S84)}\\ &- a_{xyz}xyz - a_{xyy}xy^2 - a_{xzz}xz^2 - a_{xxz}x^2z - a_{xxy}x^2y && \text{(S85)}\end{aligned}$$

If we divide $x$ into sub-species $x_1$ and $x_2$, then we have

$$\begin{aligned}\frac{\mathrm{d}x_1}{\mathrm{d}t} =& r_x x_1 - a_{xx}x_1^2 - a_{xxx}x_1^3 - a_{xy}x_1y - a_{xz}x_1z - a_{xx}x_1x_2 && \text{(S86)}\\ &- a_{xyz}x_1yz - a_{xyy}x_1y^2 - a_{xzz}x_1z^2 - a_{xxz}x_1^2z - a_{xxy}x_1^2y && \text{(S87)}\\ &- a_{xxy}x_1x_2y - a_{xxz}x_1x_2z - a_{xxx}x_1x_2^2 - a_{xxx}x_1^2x_2 && \text{(S88)}\end{aligned}$$

and

$$\begin{aligned}\frac{\mathrm{d}x_2}{\mathrm{d}t} =& r_x x_2 - a_{xx}x_2^2 - a_{xxx}x_2^3 - a_{xy}x_2y - a_{xz}x_2z - a_{xx}x_1x_2 && \text{(S89)}\\ &- a_{xyz}x_2yz - a_{xyy}x_2y^2 - a_{xzz}x_2z^2 - a_{xxz}x_2^2z - a_{xxy}x_2^2y && \text{(S90)}\\ &- a_{xxy}x_1x_2y - a_{xxz}x_1x_2z - a_{xxx}x_1x_2^2 - a_{xxx}x_1^2x_2 && \text{(S91)}\end{aligned}$$

Then

$$
\begin{aligned}
&\frac{\mathrm{d}(x_1+x_2)}{\mathrm{d}t} \\
&= r_x(x_1+x_2) - a_{xx}(x_1^2+x_2^2) - a_{xxx}(x_1^3+x_2^3) - a_{xy}(x_1+x_2)y - a_{xz}(x_1+x_2)z - 2a_{xx}x_1x_2 \\
&\quad - a_{xyz}(x_1+x_2)yz - a_{xyy}(x_1+x_2)y^2 - a_{xzz}(x_1+x_2)z^2 - a_{xxz}(x_1^2+x_2^2)z - a_{xxy}(x_1^2+x_2^2)y \\
&\quad - 2a_{xxy}x_1x_2y - 2a_{xxz}x_1x_2z - 2a_{xxx}x_1x_2^2 - 2a_{xxx}x_1^2x_2 \\
&= r_x x - a_{xx}x^2 - a_{xy}xy - a_{xz}xz - a_{xyz}xyz - a_{xyy}xy^2 - a_{xzz}xz^2 - a_{xxz}x^2z - a_{xxy}x^2y \\
&\quad - a_{xxx}(x_1^3+x_2^3+2x_1^2x_2+2x_1x_2^2)
\end{aligned} \tag{S92}
$$

In sum, label invariance dictates that the first formulation (Eqn. S82) always holds, but the second is only valid without higher-order intraspecific interactions (Eqn. S83). This subtle but fundamental distinction carries weight for deriving theoretical expectations (Bairey *et al.* 2016, Gibbs *et al.* 2022) and designing empirical tests (Mayfield & Stouffer 2017, Fox 2023).

### I Label Invariance in Stochastic Models

This appendix provides a rigorous demonstration of how the principle of label invariance constrains the form of stochastic terms in population models. We use the framework of stochastic differential equations (SDEs) to show that only noise terms with specific, mechanistically-grounded scaling properties are consistent with the principle.

#### I.1 The Label Invariance Test for SDEs

Consider a general SDE for the abundance $x_i$ of species $i$:

$$\mathrm{d}x_i = f_i(\mathbf{N})\mathrm{d}t + g_i(\mathbf{N})\mathrm{d}W_i(t)$$

Here, $f_i(\mathbf{N})$ is the deterministic drift term (e.g., population growth), which we assume has a label-invariant structure as described in the main text. The term $g_i(\mathbf{N})\mathrm{d}W_i(t)$ represents stochastic fluctuations, where $g_i$ is the noise magnitude function and $\mathrm{d}W_i$ is the increment of a Wiener process (standard Brownian motion).

The principle of label invariance requires that if we split a population $x_0$ into two ecologically identical, non-interacting subpopulations $x_1$ and $x_2$ (so that their total abundance is $x_{total} = x_1 + x_2$), the dynamics of the total population must be identical to the dynamics of the original unsplit population $x_0$.

Mathematically, let the SDE for the unsplit population be:

$$\mathrm{d}x_0 = f_0(x_0)\mathrm{d}t + g_0(x_0)\mathrm{d}W_0(t)$$

When split, the SDEs for the identical subpopulations are:

$$\begin{aligned}\mathrm{d}x_1 &= f_1(x_1, x_2)\mathrm{d}t + g_1(x_1)\mathrm{d}W_1(t)\\ \mathrm{d}x_2 &= f_2(x_1, x_2)\mathrm{d}t + g_2(x_2)\mathrm{d}W_2(t)\end{aligned}$$

By Itô's rule for addition, the dynamics of the total population are $\mathrm{d}x_{total} = \mathrm{d}x_1 + \mathrm{d}x_2$. For label invariance to hold, we must have $\mathrm{d}x_{total} \stackrel{d}{=} \mathrm{d}x_0$, where $\stackrel{d}{=}$ denotes equality in distribution. Since the deterministic part is already assumed to be invariant (i.e., $f_1 + f_2 = f_0$), this requires the stochastic parts to be equivalent:

$$g_1(x_1)\mathrm{d}W_1(t) + g_2(x_2)\mathrm{d}W_2(t) \stackrel{d}{=} g_0(x_1 + x_2)\mathrm{d}W_0(t)$$

This equality depends critically on the functional form of $g_i$ and the correlation between the noise processes $\mathrm{d}W_1$ and $\mathrm{d}W_2$.

#### I.2 Proof: Testing Common Noise Formulations

##### I.2.1 Additive Noise Violates Label Invariance

A simple case is additive noise, where the noise magnitude is a constant independent of population size: $g(N) = c$. Such a term might represent a constant, random influx or removal of individuals. Assuming the external sources are independent, the noise processes for the two subpopulations are uncorrelated.

The variance of the combined noise term for the split population is:

$$\begin{aligned}\mathrm{Var}(c\,\mathrm{d}W_1 + c\,\mathrm{d}W_2) &= \mathrm{Var}(c\,\mathrm{d}W_1) + \mathrm{Var}(c\,\mathrm{d}W_2) \quad \text{(due to independence)}\\ &= c^2\mathrm{d}t + c^2\mathrm{d}t = 2c^2\mathrm{d}t\end{aligned}$$

However, the variance of the noise for the unsplit population is $\mathrm{Var}(c\,\mathrm{d}W_0) = c^2\mathrm{d}t$. Since $2c^2\mathrm{d}t \neq c^2\mathrm{d}t$, simple additive noise is not label-invariant. Arbitrarily splitting the population artificially doubles the variance of the fluctuations.

##### I.2.2 Demographic Stochasticity Obeys Label Invariance

Demographic noise arises from the independent random fates of individuals (births, deaths).

- **Noise Magnitude:** The central limit theorem implies that the standard deviation of fluctuations scales with the square root of population size, so $g(N) = s\sqrt{N}$, where $s$ is a constant.
- **Correlation:** Because the noise stems from independent individual events, the noise processes for distinct subpopulations are **uncorrelated** (i.e., $W_1$ and $W_2$ are independent).

**Verification:** The variance of the combined stochastic term for the split population is:

$$\begin{aligned}\mathrm{Var}(s\sqrt{x_1}\mathrm{d}W_1 + s\sqrt{x_2}\mathrm{d}W_2) &= s^2\mathrm{Var}(\sqrt{x_1}\mathrm{d}W_1) + s^2\mathrm{Var}(\sqrt{x_2}\mathrm{d}W_2)\\ &= s^2(\sqrt{x_1})^2\mathrm{d}t + s^2(\sqrt{x_2})^2\mathrm{d}t = s^2(x_1+x_2)\mathrm{d}t\end{aligned}$$

This is precisely the variance of the stochastic term for the unsplit population, $s\sqrt{x_1+x_2}\mathrm{d}W_0$. Therefore, demographic noise is label-invariant.

##### I.2.3 Environmental Stochasticity Obeys Label Invariance

Environmental noise arises from external factors that affect all individuals of a given type proportionally.

- **Noise Magnitude:** The total fluctuation is proportional to population size, so $g(N) = \sigma N$, where $\sigma$ is a constant.
- **Correlation:** Because identical individuals experience the same environmental driver, their noise processes must be **perfectly correlated**, so $\mathrm{d}W_1 = \mathrm{d}W_2 = \mathrm{d}W_0$.

**Verification:** The combined stochastic term for the split population is:

$$\sigma x_1\mathrm{d}W_1 + \sigma x_2\mathrm{d}W_2 = \sigma x_1\mathrm{d}W_0 + \sigma x_2\mathrm{d}W_0 = \sigma(x_1+x_2)\mathrm{d}W_0$$

This is exactly the form of the stochastic term for the unsplit population. Therefore, environmental noise is label-invariant.

#### I.3 General Proof for Power-Law Noise

We can generalize the findings for demographic and environmental noise to show they are unique solutions for any noise magnitude of the power-law form $g(N) = cN^\alpha$. We consider a general correlation $\rho \in [-1, 1]$ between the noise processes for the two subpopulations, where $\mathrm{E}[\mathrm{d}W_1\mathrm{d}W_2] = \rho\,\mathrm{d}t$.

The variance of the combined stochastic term $cx_1^\alpha\mathrm{d}W_1 + cx_2^\alpha\mathrm{d}W_2$ is:

$$\begin{aligned}\mathrm{Var} &= \mathrm{E}\Big[(cx_1^\alpha\mathrm{d}W_1 + cx_2^\alpha\mathrm{d}W_2)^2\Big]\\ &= c^2\mathrm{E}\Big[(x_1^{2\alpha})(\mathrm{d}W_1)^2 + (x_2^{2\alpha})(\mathrm{d}W_2)^2 + 2x_1^\alpha x_2^\alpha\mathrm{d}W_1\mathrm{d}W_2\Big]\\ &= c^2\Big(x_1^{2\alpha}\mathrm{d}t + x_2^{2\alpha}\mathrm{d}t + 2\rho(x_1x_2)^\alpha\mathrm{d}t\Big)\end{aligned}$$

For label invariance, this must equal the variance of the stochastic term for the unsplit population, which is $\mathrm{Var}(c(x_1+x_2)^\alpha \mathrm{d}W_0) = c^2(x_1+x_2)^{2\alpha}\mathrm{d}t$. This gives the core identity, which must hold for all $x_1, x_2 > 0$:

$$x_1^{2\alpha} + x_2^{2\alpha} + 2\rho(x_1x_2)^\alpha = (x_1+x_2)^{2\alpha}$$

Since this must be true for any way of partitioning the population, it must hold for any ratio of $x_1$ to $x_2$. Let $x = x_1/x_2$ and divide the identity by $x_2^{2\alpha}$:

$$x^{2\alpha} + 1 + 2\rho x^\alpha = (x+1)^{2\alpha}$$

**Case 1: Uncorrelated Noise ($\rho = 0$)**

If $\rho = 0$, the identity simplifies to:

$$x^{2\alpha} + 1 = (x+1)^{2\alpha}$$

Let $h(x) = (x+1)^{2\alpha} - x^{2\alpha}$. We require $h(x) = 1$ for all $x > 0$, which means its derivative must be zero.

$$h'(x) = 2\alpha(x+1)^{2\alpha-1} - 2\alpha x^{2\alpha-1} = 2\alpha\Big[(x+1)^{2\alpha-1} - x^{2\alpha-1}\Big] = 0$$

This requires $2\alpha - 1 = 0$, which means $\alpha = 1/2$. Substituting this back confirms the result: $(x+1)^1 - x^1 = 1$. Thus, one solution is $(\alpha, \rho) = (1/2, 0)$, which corresponds to the demographic noise.

**Case 2: Correlated Noise ($\rho \neq 0$)**

If $\rho \neq 0$, we can solve the identity for $\rho$:

$$\rho(x) = \frac{(x+1)^{2\alpha} - x^{2\alpha} - 1}{2x^\alpha}$$

For $\rho$ to be a constant, its derivative with respect to $x$ must be zero for all $x > 0$. Setting the numerator of $\mathrm{d}\rho(x)/\mathrm{d}x$ to zero gives:

$$\alpha\Big[(x-1)(x+1)^{2\alpha} - (x+1)(x^{2\alpha} - 1)\Big] = 0$$

This equation holds if $\alpha = 0$. However, substituting $\alpha = 0$ back into the expression for $\rho(x)$ gives $\rho = 0$, which contradicts our assumption that $\rho \neq 0$.

Alternatively, the term in the square brackets must be zero. The only solution is $\alpha = 1$. Substituting $\alpha = 1$ into the expression for $\rho(x)$ yields $\rho = 1$.

Therefore, the only solution for the correlated noise case is $(\alpha = 1, \rho = 1)$, which corresponds to environmental noise.

#### I.4 Itô vs. Stratonovich Interpretation

A final consideration is the choice of stochastic calculus. The Stratonovich interpretation of an SDE, $\mathrm{d}N = f\mathrm{d}t + g \circ \mathrm{d}W$, is equivalent to an Itô SDE with an added "noise-induced drift" term:

$$\mathrm{d}N = \Big(f + \frac{1}{2}g'g\Big)\mathrm{d}t + g\,\mathrm{d}W$$

We must check if this drift term preserves label invariance.

- **For environmental noise** ($g = \sigma N$), the drift term is $\frac{1}{2}(\sigma)(\sigma N) = \frac{1}{2}\sigma^2 N$. Since this term scales linearly with $N$, it is label-invariant.

- **For demographic noise** ($g = s\sqrt{N}$), the drift term is $\frac{1}{2}(\frac{1}{2}sN^{-1/2})(s\sqrt{N}) = \frac{1}{4}s^2$. This is a constant. If we split population $x_0$, the total drift for $x_1 + x_2$ would be $(\frac{1}{4}s^2 + \frac{1}{4}s^2) = \frac{1}{2}s^2$, which does not match the drift for the unsplit case. Thus, a Stratonovich interpretation of demographic noise **violates label invariance**.

Fortunately, this is not a practical concern, as demographic stochasticity is typically derived from first principles (e.g., via the chemical Langevin approximation of a master equation), which naturally leads to the Itô formulation without a noise-induced drift. For environmental noise, both interpretations can be valid, but the final Itô form used for analysis must be checked for invariance.
